# Multi-omics integration reveals sex-based differences in the circulating extracellular vesicle lipidome and miRNome of alcohol use disorder patients

**DOI:** 10.1101/2025.07.15.664861

**Authors:** Carla Perpiñá-Clérigues, Susana Mellado, Cristina Galiana-Roselló, Saritha Kodikara, Blanca Martín-Urdiales, Miguel Marcos, Kim-Anh Lê Cao, Francisco García-García, María Pascual

**Affiliations:** Department of Physiology, School of Medicine and Dentistry, University of Valencia, 46010 Valencia, Spain; Computational Biomedicine Laboratory, Príncipe Felipe Research Center, 46012 Valencia, Spain; Melbourne Integrative Genomics, School of Mathematics and Statistics, University of Melbourne, Melbourne, VIC, Australia; Department of Internal Medicine, University Hospital of Salamanca, University of Salamanca, Institute of Biomedical Research of Salamanca (IBSAL), 37007 Salamanca, Spain

**Keywords:** lipidomics, miRNA transcriptomics, multi-omics, extracellular vesicles, alcohol use disorder, sex-based differences

## Abstract

Integrated multi-omics and extracellular vesicle (EV) analysis are emerging as powerful, complementary strategies for biomarker discovery. These approaches offer promising tools to enhance early detection, diagnosis, and treatment of alcohol use disorder (AUD). Here we applied an integrated miRNomic and lipidomic approach to analyze plasma EVs from AUD patients and controls of both sexes to gain a comprehensive understanding of the underlying molecular mechanisms. We identified an AUD signature with predictive potential for diagnostic applications. Individual features (e.g., hsa-miR-99b-3p, hsa-miR-556-5p, Cer_NDS-d39:1, and PI18:0_18:2) represented important components; however, the strength of this signature lay in the combined profile rather than isolated markers. We also revealed an AUD-sex signature that provided insight into how biological responses to alcohol differ between females and males (including features such as hsa-miR-1301-3p and PC39:4), which also underscored the power of multi-omic integration. The individual miRNome approach also revealed an opposite functional alteration by sex in various alcohol related systems, such as pathways associated with immunity, oxidative stress, and autophagy. An open-access Shiny web application (https://carpercle.shinyapps.io/SexEVEthOmics/) accompanies this study, providing interactive access to the complete dataset and additional analyses for customized exploration. Together, our findings underscore the added value of multi-omics integration in identifying clinically relevant molecular signatures of disease; this sex-informed approach offers a promising path toward more personalized diagnostic tools and therapeutic strategies in AUD.

## Introduction

Multi-omics has emerged as a result of advances in various omic-based technologies (including genomics and metabolomics) alongside enhanced computing capabilities, which have enabled an improved understanding of the complex molecular interplay between health and disease by combining the power of individual data types [1]. This analytical approach measures different layers of cellular information, facilitating the identification of molecular signatures and biomarkers for mechanism-based classifications and tailored therapeutic interventions [2,3] and offering deep insight into the pathophysiology of disease states [4]. The use of multivariate approaches - facilitated by this novel integrative analysis - has enabled the expansion of results obtained through univariate methods [5] to predict, prevent, and treat disease more precisely by considering each individual’s genetics, environment, and lifestyle factors [1].

Recent reports support the applicability of transcriptomics and lipidomics in biomarker identification and pathology elucidation. Transcriptomics, including miRNA analysis, represents a robust methodology for determining mRNA/miRNA expression via high-throughput RNA-sequencing (RNA-seq) [4]. Although lipidomics represents an effective approach to analyzing changes to the levels of endogenous lipids in biological systems [4], alterations in lipid levels/enzymatic reactions may not accurately identify pathways altered in specific pathologies. We can address this gap by integrating various omic-based technologies, which offer a more comprehensive view of alterations in complex biological processes induced by any pathological condition. A combined miRNomic and lipidomic approach has the potential to generate extensive datasets that will support the identification of associations between miRNAs and lipids/enzymes and uncover molecular mechanisms based on high-throughput data.

An increasing number of studies into alcohol consumption have applied transcriptomic and lipidomic technologies to understand the molecular mechanisms underlying alcohol use disorders (AUDs) and search for possible disease biomarkers [6–8]. Alcohol remains a widely misused substance, with the World Health Organization reporting that abuse causes approximately 2.6 million deaths per year [9]. Early AUD identification and treatment potentially reduce the risk of developing related pathologies (e.g., certain types of cancer, diseases associated with reduced immunity, and increased infections, infertility, and depression) [10]. Overall, the lack of specific and sensitive biomarkers for AUD diagnosis and effective treatment approaches has prompted the need for the application of novel technologies such as integrated multi-omics. Multi-omics enables the simultaneous analysis of miRNAs and lipids, for example, to provide insights into miRNomic and lipidomic alterations following alcohol consumption, which will help to map complex relationships.

Extracellular vesicles (EVs), which carry miRNAs and lipids, act as critical intercellular mediators, particularly under cellular stress or damage. Unlike synthetic drug delivery systems, EVs demonstrate superior biocompatibility, extended biodistribution, and low immunogenicity [11]. These properties position EVs and their content as valuable non-invasive biomarkers and therapeutic targets for diverse pathologies, including neurodegenerative disorders [12–14]. We previously identified AUD-related lipidomic fingerprints in plasma EVs from AUD females and males [6]; here, the first implementation of an integrated multi-omics approach (combining transcriptomics and lipidomics) using plasma EVs isolated from AUD males and females has enabled a more comprehensive understanding of the biological processes involved following excessive alcohol consumption.

## Materials and Methods

### Human Subjects and Experimental Design

This study includes AUD patients (according to DSM-5 criteria) referred to the Alcoholism Unit of the University Hospital of Salamanca (Spain) [15]. Healthy volunteers who reported drinking < 15 g of ethanol/day were also recruited as control patients. AUD patients and healthy volunteers were previously recruited and described in Perpiñá-Clerigués *et al*. 2024 [6]. All patients in this group actively drank ≥ 90 g of ethanol/day until entering the study. Sample numbers per experimental group varied across omics (see Table S1 for details). All individuals gave their informed consent to participate, and the study was approved by the Ethics Committee of the University Hospital of Salamanca (Spain). Plasma samples were snap-frozen in liquid nitrogen and stored at -80 °C until further use. At this point, samples were processed for EV isolation.

Investigating the effects of alcohol consumption on miRNA and lipid content in EVs and exploring the molecular basis underlying sex-specific responses used the strategy depicted in Figure 1. Lipids and miRNAs were obtained from plasma EVs of AUD and control patients.

**Figure 1.**
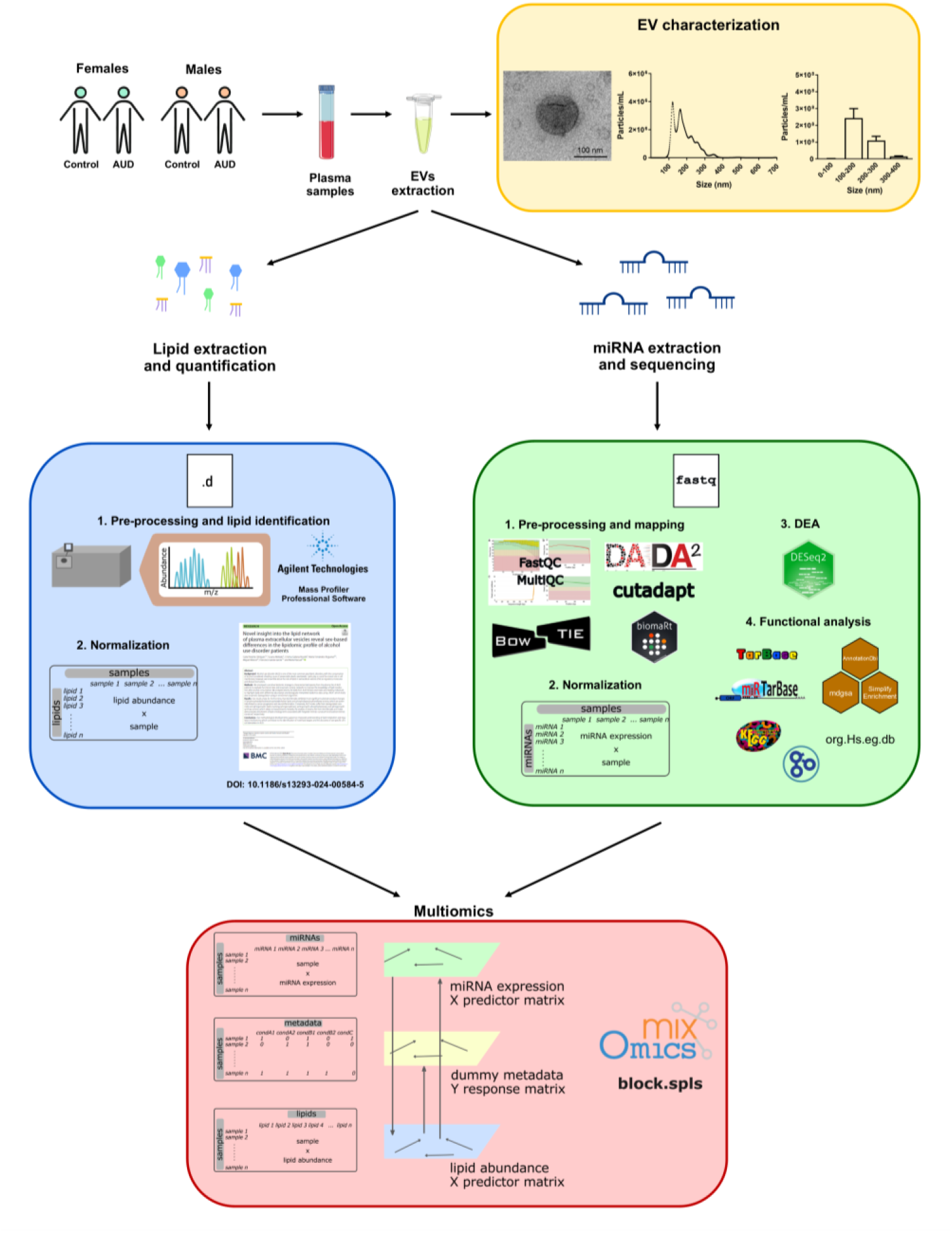
Experimental design and workflow. Lipids and miRNAs were extracted from plasma extracellular vesicles (EVs) of alcohol use disorder (AUD) patients and healthy individuals of both sexes. The lipidome and miRNome were analyzed independently before the data were integrated to examine reciprocal regulation. The lipidome analysis was previously conducted by Perpiñá-Clérigues *et al*. [6], who detailed the workflow from lipid extraction to bioinformatics characterization of EV lipid content across experimental groups. The miRNome analysis involved four key steps: (i) extraction and sequencing of EV miRNAs; (ii) preprocessing, mapping, and normalization; (iii) differential expression analysis; and (iv) functional profiling. The integration of miRNomic and lipidomic datasets begins with the normalization of expression matrices. The implementation of the mixOmics R package applied the block sparse partial least squares (sPLS) method for multi-omics integration.

### EV Isolation and Characterization

Human plasma EVs were isolated and characterized by transmission electron microscopy (TEM) and nanoparticle tracking analysis (NTA) (Figure 1), as previously described by Ibáñez *et al*. [16].

### RNA Extraction and Sequencing

Total RNA from plasma EVs was isolated using the Total Exosome RNA Isolation Kit (Invitrogen, Waltham, MA, USA), following the manufacturer’s instructions. The construction of miRNA libraries was performed using the TruSeq Small RNA Library Preparation Kit from Illumina (San Diego, CA, USA) at the Genomics and Genetics Service of the Príncipe Felipe Research Center (Valencia, Spain). RNA transcripts were first converted into complementary (c)DNA libraries, followed by quality and quantity assessments employing High Sensitivity D1000 ScreenTape for TapeStation System (Agilent Technologies, Santa Clara, CA, USA). The final libraries were pooled in equimolar concentration for sequencing in the MiSeq™ System (Illumina) through MiSeq™ Reagent Nano Kit v2 (Illumina) with 1 x 50 bp read length. Next-generation sequencing was performed at the Centre for Genomic Regulation (CRG) in Barcelona (Spain). The raw data generated by the high-throughput sequencing of miRNA were exported as fastq files.

To validate the miRNA levels from plasma EVs, real-time quantitative PCR (RT-qPCR) was performed as previously described [16]. Details of the nucleotide sequences of the used miRNA assays can be found in the Supplementary Material (Table S2).

### Processing and Mapping

Raw sequencing reads underwent preprocessing using Cutadapt (v.4.6) [17]. Sequences between 17– 35 nucleotides and with Phred scores ≥30 were retained. Illumina adapters were removed before downstream analysis. Trimmed reads were mapped to human mature miRNAs from miRBase (mature.fa) [18] using Bowtie2 (v2.5.3) [19]. The alignment parameters (-L 6 -i S,0,0.5 --ignore-quals -- norc --score-min L,-1,-0.6 -D 20 --very-sensitive) were adapted from Locati et al. [20], optimizing sensitivity and specificity. Read counts were extracted from the SAM files, and only uniquely mapped reads—filtered via Bash—were used for quantification to avoid biases from multimapping.

### Exploratory Analysis and Normalization of miRNA Profiles

Custom R scripts utilizing DESeq2 (v.1.42.1) [21] were used for both filtering and normalization. miRNAs ≥10 reads in at least three samples were retained. Subsequently, the median ratio normalization method implemented in DESeq2 was applied to adjust for differences in sequencing depth and RNA composition. Boxplots, PCA, and clustering—performed pre- and post-normalization—assessed group distribution, batch effects, and data quality. All statistical analyses and data visualization scripts were executed in R (v4.3.2) [22] using RStudio as the integrated development environment.

### Differential Expression Analysis of miRNAs

Identifying differentially expressed miRNAs employed DESeq2 [21]. P-values were adjusted for multiple testing using the Benjamini–Hochberg (BH) method [23], with statistically significant miRNAs defined as those with a BH-adjusted p-value ≤0.05.

Three primary comparisons were performed to assess sex-specific effects of AUD:

1. AUD Impact in Females (**IF**) assesses the differences between AUD females and control females (AUD.Females - Control.Females)
2. AUD Impact in Males (**IM**) assesses the differences between AUD males and control males (AUD.Males - Control.Males)
3. Impact of Sex in AUD (**IS**) assesses the differences between IF and IM (AUD.Females - Control.Females) - (AUD.Males - Control.Males)

Expression changes were quantified using log fold change (LFC), where the absolute value represents the magnitude of change, and the sign indicates direction. A positive LFC indicates higher expression in the first group (AUD groups for IF/IM; females for IS), while a negative LFC indicates higher expression in the second group (controls for IF/IM; males for IS). For the IS comparison, which examines sex-dependent responses to AUD, the interpretation requires careful consideration. To resolve these scenarios, individual miRNA trends in IF and IM must be cross-referenced (Figure S1, “Help” section of our website https://carpercle.shinyapps.io/SexEVEthOmics)

A complementary analysis examined combined AUD versus control samples (without sex stratification) to assess whether sex-specific analysis provided additional insights (see Supplementary Material).

### Functional Profiling of miRNAs

Differentially expressed miRNAs were linked to their experimentally validated target genes using combined data from TarBase [24] and miRTarBase [25]. Next, their LFC and p-values were transferred to target genes using the mdgsa package (v.0.99.2) [26], reflecting both statistical significance and direction of regulation. This allowed modeling of the cumulative regulatory effect of multiple miRNAs on individual genes. Functional enrichment analysis of Gene Ontology (GO) terms (27) and Kyoto Encyclopedia of Genes and Genomes (KEGG) pathways (28) was performed using gene annotations from org.Hs.eg.db R package (v. 3.18.0) [27], with significance determined by BH-adjusted p-value ≤0.05. Pathways with divergent regulatory impacts between sexes were identified based on opposing log odds ratios (LORs), where the sign and magnitude of the LOR reflected the direction and strength of the enrichment effect. Visualization was carried out using simplifyEnrichment [28] (semantic similarity threshold = 0.8) and dot plots to illustrate functional divergence across conditions.

### Multi-omic Integration

Data were analyzed using processed and normalized results from previous univariate analyses. To assess concordance between datasets, an initial sPLS analysis was performed on miRNA and lipid data, evaluating their alignment with AUD and sex conditions. Then, a multi-omic integration was then conducted using the N-integration approach via the block.sPLS method (mixOmics v.6.26.0) [5], with miRNA and lipid levels as predictors (X) and a dummy matrix encoding AUD-sex groups as response (Y). The model was fitted using three components with a fully connected design between blocks (design = “full”). Feature selection retained 20, 20, and 10 features in X, and 6, 6, and 4 in Y for components 1 to 3, respectively. Variable contributions and inter-dataset relationships were explored using plotVar and plotLoadings.

### Web Platform - SexEVEthOmics

In addition to the results presented in this article, an open-access, interactive web platform (https://carpercle.shinyapps.io/SexEVEthOmics/) was developed using the R shiny framework (v.1.10) [29]. This tool enables users to access the full dataset and interactively explore differential miRNA expression results, functional analyses, lipidomic profiles, and multi-omics integration. The platform is structured into six sections and provides access to additional analyses and visualizations not included in the main manuscript. Some modules allow users to adjust parameters, offering opportunities for customized exploration and hypothesis generation.

## Results

### Sex-based Differences in the miRNA Expression Profile of AUD Patient Plasma EVs

To implement our knowledge in the search for biomarkers in plasma EVs from AUD patients, we analyzed plasma EV samples from female and male AUD and healthy patients (controls) who had been previously assessed via lipidomics [6]. We investigated the role of miRNAs in the four experimental groups, defined by sex and chronic alcohol consumption. Figure 2A-B reports the upregulated (LFC > 0) and downregulated (LFC < 0) miRNAs. We observed 4 upregulated and 3 downregulated miRNAs in females and 18 upregulated and 4 downregulated miRNAs in males. Notably, the comparison in males revealed a higher number of upregulated miRNAs compared to females (Figure 2A-B), suggesting distinct sex-specific responses to alcohol consumption. Notably, we failed to detect any significantly altered miRNAs in the IS comparison (Web Platform - Study Overview [https://carper-cle.shinyapps.io/SexEVEthOmics/], Table S3).

**Figure 2.**
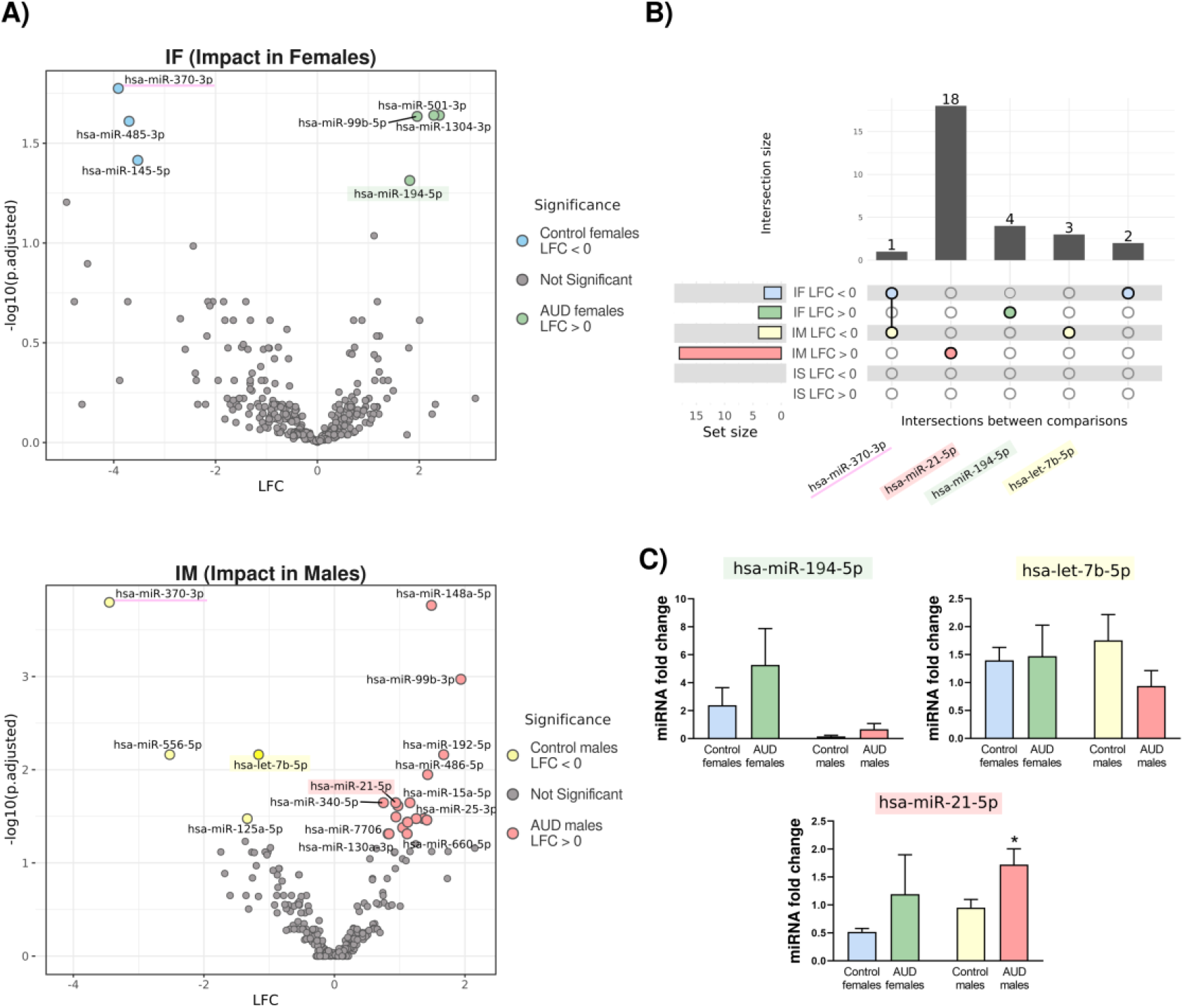
Differential Expression and Validation of miRNAs in Plasma EVs from AUD and Control Patients Based on Sex. **A)** Volcano plots demonstrate differential miRNA expression (adjusted p-value < 0.05) in the IF and IM comparisons. miRNAs with a positive log fold change (LFC > 0) display upregulation (green and red dots for IF and IM, respectively) and miRNAs with a negative LFC (LFC < 0) display downregulation (blue and yellow dots for IF and IM, respectively) in AUD patients. Highlighted miRNAs were validated by RT-qPCR, and underlined miRNA (hsa-miR-370-3p) represents common significantly altered miRNA between sexes. **B)** Intersection plot displaying shared and unique significantly altered miRNAs across IF, IM, and IS comparisons. miRNAs with names under the dots belong to that intersection pattern. **C)** RT-qPCR validation of three selected miRNAs: *hsa-miR-194-5p, hsa-let-7b-5p*, and *hsa-miR-21-5p*. Data represent mean ± SEM, n = 4-6 samples/group. * p < 0.05, compared to healthy controls. One-way ANOVA with Tukey multiple comparisons testing was performed. *AUD: alcohol use disorder, IF: AUD Impact in Females, IM: AUD Impact in Males, IS: Impact of Sex in AUD*.

We next validated miRNA-seq data using RT-qPCR (Figure 2C). We selected three miRNAs (hsa-miR-194-5p, hsa-let-7b-5p, and hsa-miR-21-5p) involved in the immune response [30–33], which revealed significant changes in AUD patients. Consistent with the miRNA-seq results, hsa-miR-194-5p and hsa-miR-21-5p tended to increase in AUD females compared to healthy females, while hsa-let-7b-5p tended to decrease and hsa-miR-21-5p significantly increased in AUD males compared to healthy males. Overall, the data suggest that these miRNAs may participate in AUD pathogenesis in a sex-dependent manner.

### Sex-based Differences in miRNA Functional Analysis of Plasma EVs from AUD Patients

We next assessed the functional impact of miRNA deregulation in AUD patient plasma EVs to provide biological insights into associated etiopathogenic pathways [34]. We performed gene set analysis (GSA) on validated target genes associated with the differentially expressed miRNAs, incorporating GO terms related to Biological Processes (BP), Cellular Components (CC), and Molecular Functions (MF), and KEGG pathways. Notably, we observed several significantly impacted GO terms and KEGG pathways across IF, IM, and IS comparisons, highlighting notable sex-based differences. Table S4 summarizes the number of significant GSA results across the IF, IM, and IS comparisons, while Figures S2–S5 display the common significant results between IF and IM for the BP, CC, and MF ontologies. Figure 3A presents clusters derived from 66 common BP GO terms that exhibit significant sex-specific variation across all comparisons, characterized by negative LOR values in females and positive LOR values in males. These processes cluster into four main functional groups: (1) gene expression and protein turnover, encompassing ubiquitin-dependent degradation, mRNA regulation, splicing, phosphorylation, and catabolic pathways; (2) cell signaling and fate determination, such as cellular transformation, apoptosis, growth regulation, and receptor-mediated signaling; (3) cellular structure and stress response; and (4) vesicle-mediated transport related to intracellular trafficking and membrane dynamics.

**Figure 3.**
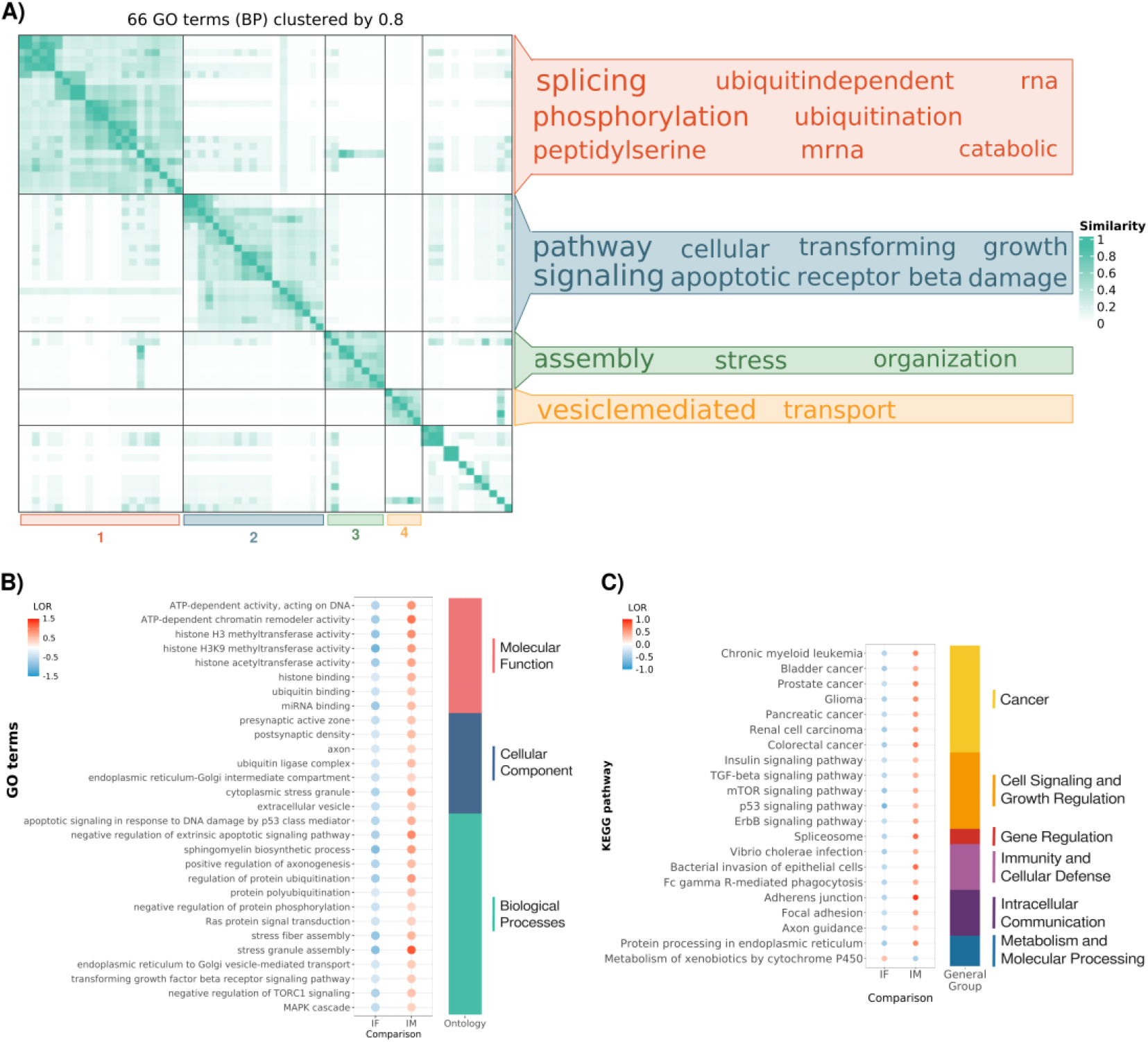
Sex-Specific Functional Signatures of Plasma EV miRNAs in AUD Patients - Gene Set Analysis Results. **A)** Clustering of significant BP GO terms common across IF, IM, and IS comparisons based on semantic similarity (threshold ≥0.8). Word clouds summarize representative terms for each cluster. **B)** Dot plot depicting selected significant GO terms (absolute LOR > 0.3) across MF, CC, and BP ontologies. Dot color represents the LOR value, and the right-side color bars indicate the ontology category of each term. **C)** Dot plot illustrating significant KEGG pathways (absolute LOR > 0.3). Dot color represents the LOR value, while the right-side color bars classify pathways into general functional groups. *AUD: alcohol use disorder, BP: Biological Processes, CC: Cellular components, IF: AUD Impact in Females, IM: AUD Impact in Males, GO: Gene Ontology, KEGG: Kyoto Encyclopedia of Genes and Genomes, LOR: log odds ratio, MF: Molecular Functions*.

Figure 3B illustrates a selection of the most relevant significantly altered GO terms characterized by distinct sex-dependent patterns.

The MF ontology highlighted terms related to histone modifications, including histone H3 methylation and ubiquitin-related processes. The CC ontology revealed an enrichment in terms related to synaptic activity, including the pre- and post-synaptic regions, the axon, and EVs. Within the BP ontology, significant terms associated with neurological functions (e.g., axonogenesis and lipid metabolism), cellular communication, and vesicle-mediated transport, including Golgi-related pathways, Target of Rapamycin Complex 1 (TORC1) signaling, protein phosphorylation, and Mitogen-Activated Protein Kinase (MAPK) cascades. Interestingly, only terms associated with DNA replication - specifically, the BP term “regulation of the DNA damage checkpoint” and the CC term “CMG complex” -exhibited a consistent pattern across IF and IM comparisons (Figures S2– S4).

Finally, KEGG pathway enrichment analysis revealed divergence between sexes in the functional impact of miRNA deregulation in AUD (Figure 3C). In AUD females (IF), several pathways displayed a negative LOR, whereas these pathways possessed a positive LOR in AUD males (IM). These pathways participate in a range of disease-related processes, including cancer-associated pathways linked to the excretory and digestive systems (e.g., renal cell carcinoma, bladder, pancreatic, and colorectal cancer), pathways related to cell signaling and growth regulation (e.g., mammalian target of rapamycin [mTOR], p53, and the family of receptor tyrosine kinases ErbB), immunity-related pathways, intracellular communication (e.g., axon guidance and adherens junctions), and metabolism and molecular processing associated with the endoplasmic reticulum.

### Integrated Transcriptomic and Lipidomic Analyses Reveal Sex-based Differences in AUD Patient Plasma EVs

We employed our miRNomic data, combined with lipidomics data from [6], to explore novel relationships between miRNAs and lipids in plasma EVs isolated from AUD females and males. After data preprocessing and normalization according to their respective omic natures, we obtained a total of 387 miRNAs and 575 lipids for the same samples. Figure S6 illustrates the sPLS results, where the samples exhibit comparable spatial organization in both datasets, with high correlation values observed for the first two components (Comp1: 0.951 and Comp2: 0.953). This strong alignment suggests the successful application of the N-integration methodology (block.sPLS).

### Biological Patterns Captured by Components 1 and 2: Links to Alcohol Consumption and Sex

Performing dimension reduction using the mixOmics package produces latent components defined by their loading vectors, which indicate the weight of each original variable’s contribution to the component. Figure 4 depicts the sample plot and the loadings plot for Components 1 and 2. Component 1 captures variance related to alcohol consumption, while Component 2 captures variance related to sex. The bar plots display the loading values, indicating the extent to which each feature contributes to the differentiation of the groups; those at the bottom and those with longer bars have the most significant influence. As a result, we identified a subset of 40 miRNAs (Figure 4B) and 40 lipids (Figure 4C) that may distinguish and clarify the influence of alcohol (e.g., hsa-miR-148a-5p, hsa-miR-7706, and hsa-miR-21-5p and Cer_NDS d39:1_neg, EtherPC 16:0e_18:2_neg, and SM d18:2_23:0) and sex (e.g., hsa-miR-30d-5p, hsa-miR-152-3p, and hsa-miR-1271-5p and Cer_NDS d39:1_neg, EtherPC 16:0e_18:2_neg, and SM d18:2_23:0). Table S5 details the description of lipid subclass abbreviations.

**Figure 4.**
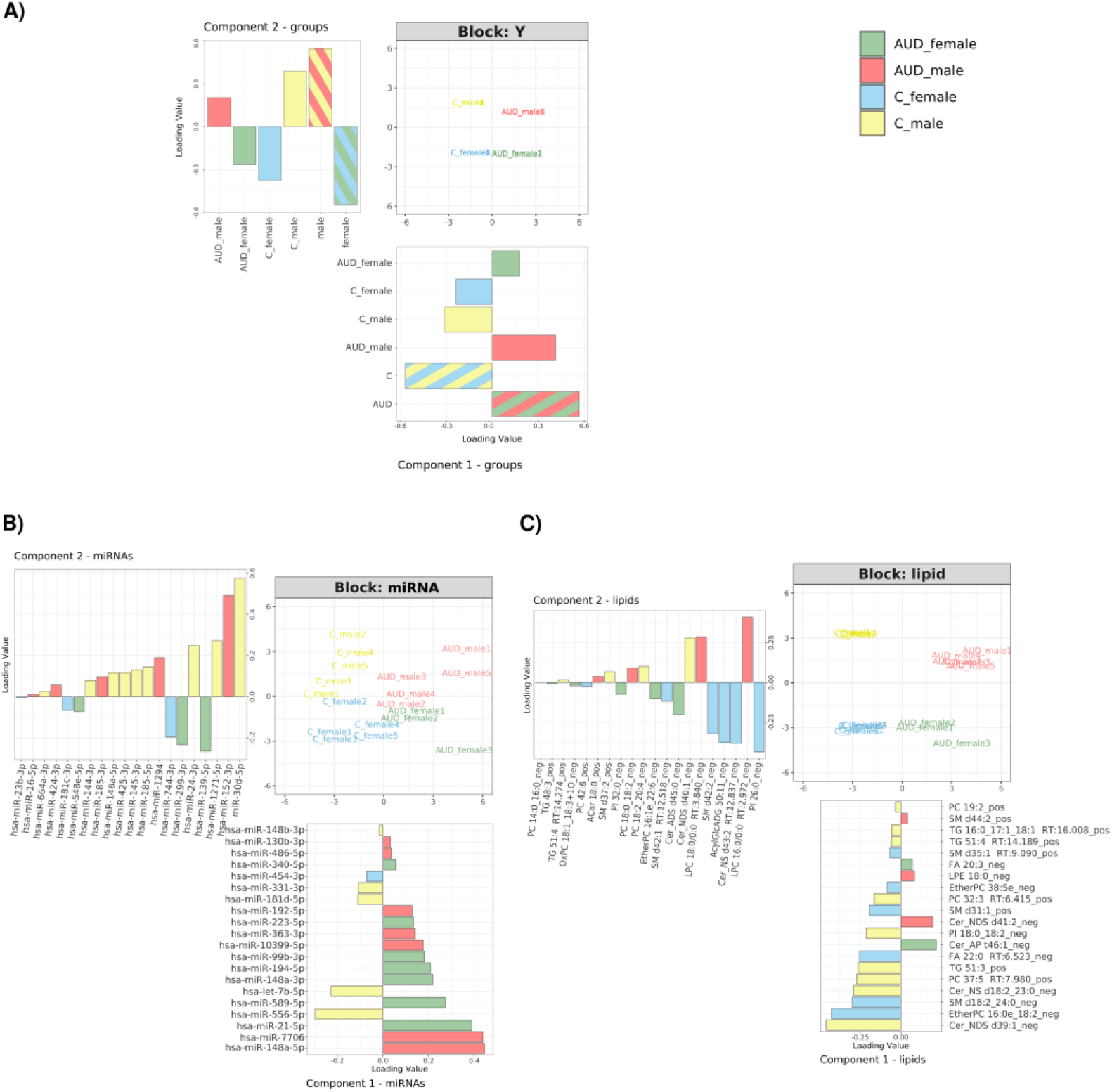
AUD- and Sex-driven Sample Separation Based on Integrated Patient EV miRNA and Lipid Profiles. Sample and loading plots from the block.sPLS integration analysis performed using mixOmics, representing Components 1 and 2. Samples are projected into the space defined by the components derived from the **(A)** metadata (Y), **(B)** miRNA, and **(C)** lipid dataset. The loading plots identify the most relevant variables contributing to each component: **(A)** metadata (Y), **(B)** miRNA, and **(C)** lipids. Component 1 is represented on the X-axis, with its corresponding loading plot displayed behind the sample plot. Component 2 is shown on the Y-axis, with its loading plot displayed to the left. Colors in the sample plots indicate the experimental groups. Colors in the loading plots denote which experimental group has the highest mean expression/abundance of that feature.

A correlation plot with a cutoff of 0.5 illustrated the relationship between the miRNomic and lip-idomic datasets and the experimental groups (Figure 5). In line with our focus on integrating these molecular features into a comprehensive physiological and pathological framework, Figure 5 highlights those lipids and miRNAs associated with pathologies/terms related to alcohol consumption (Table S6 provides more detailed information). The features positioned in the upper and lower areas associate with males and females, respectively, while the features on the right and left sides correspond to AUD and control patients, respectively.

**Figure 5.**
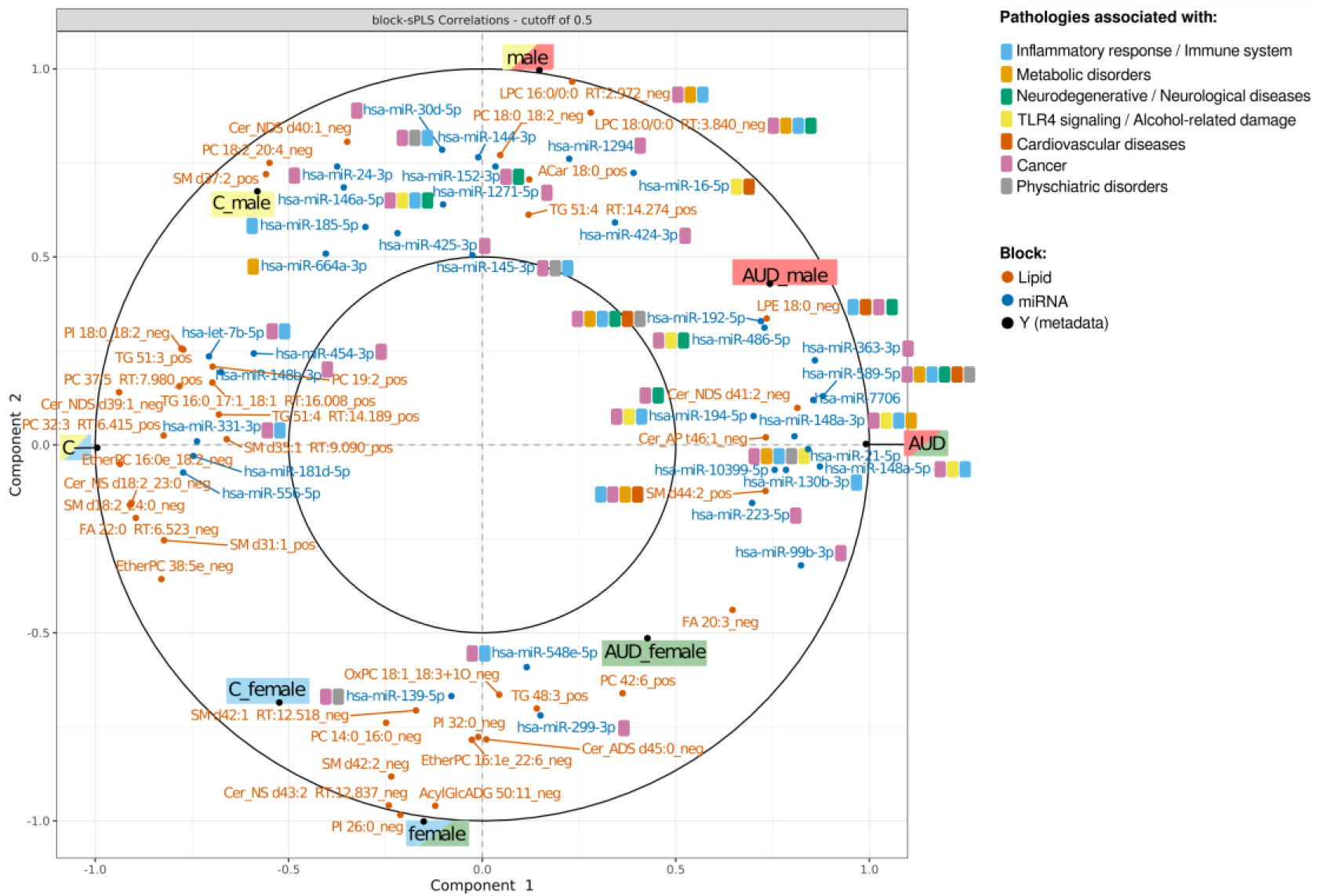
Varplot from the block.sPLS analysis displaying Component 1 vs. Component 2 (cutoff = 0.5). The plot displays the correlations between selected variables from the miRNA and lipidomic blocks and the experimental groups (Y block), projected onto the first two components. Only variables with a correlation ≥ |0.5| are shown. Labels are colored by block (lipid, miRNA, or Y), and icons indicate the associated pathologies based on literature annotation. Sample group centroids are highlighted (e.g., AUD female, control male), and distribution patterns reveal distinct molecular signatures by sex and alcohol exposure.

Table 1 presents the AUD signature of 20 miR-NAs and 20 lipids identified through the integrated joint model that maximizes the separation between AUD patients and controls. This multivariate panel effectively captured the combined discriminative power of both molecular layers; additionally, by integrating results from univariate linear regression analyses (DEA: Differential expression analysis of miRNome data, DAA: Differential abundance analysis of lipidome data), we highlighted the miRNAs and lipids that individually differentiate the experimental groups in at least one sex. We consider that these features contribute additional robustness to the panel, representing potential biomarkers with strong discriminative capacity (relevant in a sex-specific context).

**Table 1.** AUD Signature Based on Multi-omics Integration (Component 1).

| FEATURE | PATTERN | DEA/<br>DAA | FEATURE | PATTERN | DEA/<br>DAA |
| --- | --- | --- | --- | --- | --- |
| Cer_AP t46:1_neg | + | F | Cer_NS d18:2_23:0_neg | - | F & M |
| Cer_NDS d41:2_neg | + |  | EtherPC 16:0e_18:2_neg | - | F & M |
| FA 20:3_neg | + | M | EtherPC 38:5e_neg | - | M |
| LPE 18:0_neg | + |  | FA 22:0 RT:6.523_neg | - |  |
| SM d44:2_pos | + |  | PC 19:2_pos | - | M |
| hsa-miR-10399-5p | + | M | PC 32:3 RT:6.415_pos | - |  |
| hsa-miR-130b-3p | + |  | PC 37:5 RT:7.980_pos | - |  |
| hsa-miR-148a-3p | + | M | PI 18:0_18:2_neg | - | M |
| hsa-miR-148a-5p | + | M | SM d18:2_24:0_neg | - | F & M |
| hsa-miR-192-5p | + | M | SM d31:1_pos | - |  |
| hsa-miR-194-5p | + | F | SM d35:1 RT:9.090_pos | - |  |
| hsa-miR-21-5p | + | M | TG 16:0_17:1_18:1 RT:16.008_pos | - |  |
| hsa-miR-223-5p | + |  | TG 51:3_pos | - |  |
| hsa-miR-340-5p | + | M | TG 51:4 RT:14.189_pos | - |  |
| hsa-miR-363-3p | + | M | hsa-let-7b-5p | - | M |
| hsa-miR-486-5p | + | M | hsa-miR-148b-3p | - |  |
| hsa-miR-589-5p | + |  | hsa-miR-181d-5p | - |  |
| hsa-miR-7706 | + | M | hsa-miR-331-3p | - |  |
| hsa-miR-99b-3p | + | M | hsa-miR-454-3p | - |  |
|  |  |  | hsa-miR-556-5p | - | M |

### Divergent Patterns Based on AUD and Sex Captured by Component 3

Figure 6A displays the loading plots for Component 3, highlighting opposing patterns between the main experimental groups and a set of 10 miRNAs and 10 lipids that explain the variance in this component (AUD-sex signature). Figure 6B-C presents the mean values of these variables to visualize miRNA expression and lipid abundance patterns across the experimental groups. Three main patterns emerge from these results: (I) the pattern in light gray indicates an increase in AUD females and a decrease in AUD males compared to controls (e.g., hsa-miR-1301-3p and Cer_NS d18:1_24:1) (Figure S1-A1); (II) the pattern in dark gray indicates a decrease in AUD females and an increase in AUD males relative to controls (e.g., hsa-miR-1260b and Cer_NS d18:1_22:0) (Figure S1-B6); and (III) the pattern in white indicates stability when comparing controls of both sexes, with a specific variation (increase or decrease) in one sex within AUD patients relative to its corresponding control (e.g., hsa-miR-197-3p) (Figure S1-A2, -B7, and B9). Patterns I and II exhibit an apparent cross-over effect, as the response to AUD follows an opposing trend between sexes, with increases in one sex corresponding to decreases in the other. These patterns suggest a sex-dependent response to AUD, where specific miRNAs and lipids exhibit opposite expression trends between males and females. Notably, we found that specific lipids identified in these patterns displayed significance or a trend towards significance in the individual analysis for the IS comparison [6], reinforcing the impact of sex on the impact of AUD. In contrast, the transcriptomics analysis failed to reveal significant results for the IS comparison (Figure 2); however, a set of miRNAs emerged through the integrative approach, highlighting the influence of sex in AUD. Table 2 summarizes the AUD-sex signature.

**Table 2.**
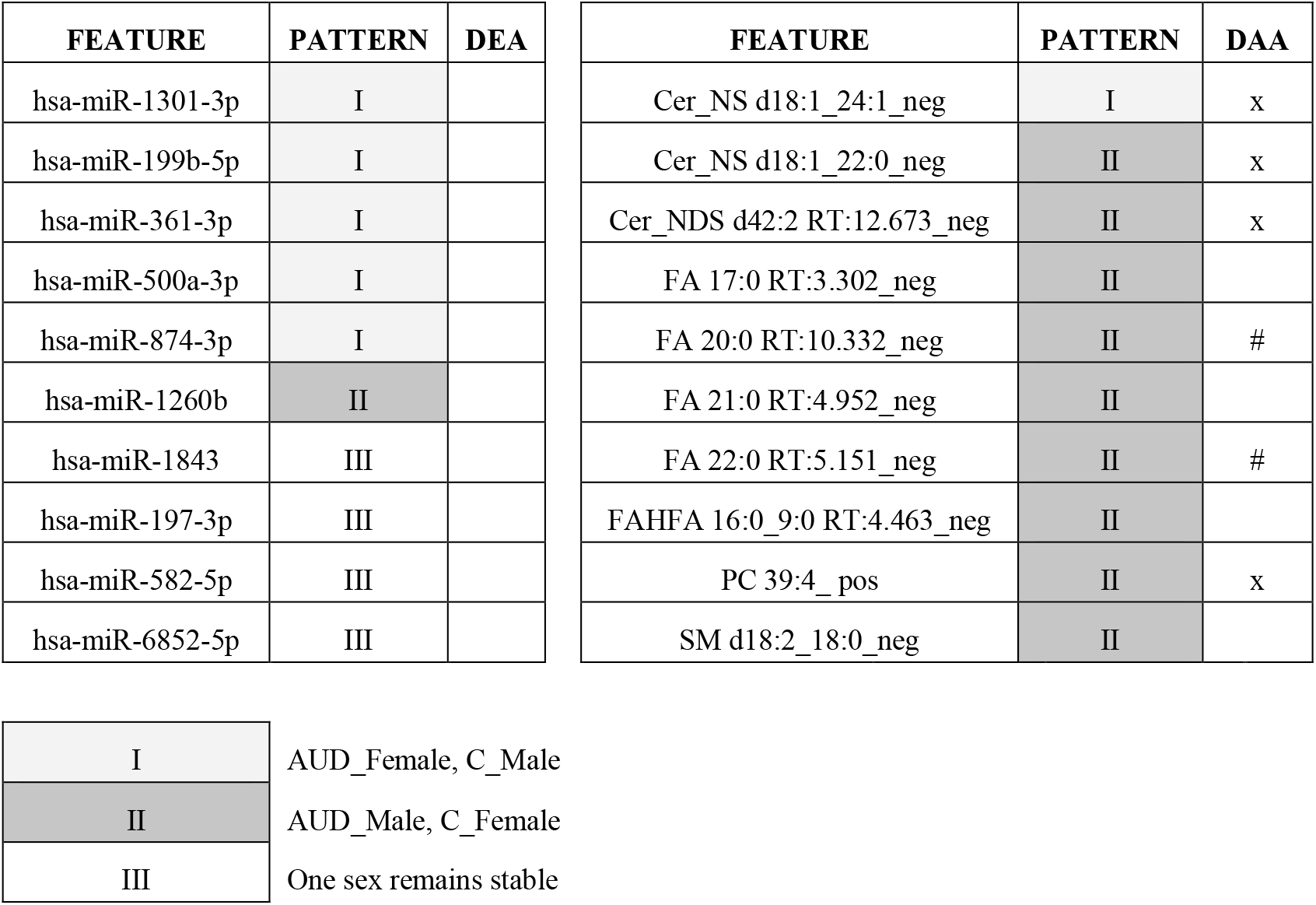
AUD-Sex Signature Based on Multi-omics Integration (Component 3).

| FEATURE | PATTERN | DEA | FEATURE | PATTERN | DAA |
| --- | --- | --- | --- | --- | --- |
| hsa-miR-1301-3p | I |  | Cer_NS d18:1_24:1_neg | I | x |
| hsa-miR-199b-5p | I |  | Cer_NS d18:1_22:0_neg | II | x |
| hsa-miR-361-3p | I |  | Cer_NDS d42:2 RT:12.673_neg | II | x |
| hsa-miR-500a-3p | I |  | FA 17:0 RT:3.302_neg | II |  |
| hsa-miR-874-3p | I |  | FA 20:0 RT:10.332_neg | II | # |
| hsa-miR-1260b | II |  | FA 21:0 RT:4.952_neg | II |  |
| hsa-miR-1843 | III |  | FA 22:0 RT:5.151_neg | II | # |
| hsa-miR-197-3p | III |  | FAHFA 16:0_9:0 RT:4.463_neg | II |  |
| hsa-miR-582-5p | III |  | PC 39:4_pos | II | x |
| hsa-miR-6852-5p | III |  | SM d18:2_18:0_neg | II |  |

|  |  |
| --- | --- |
| I | AUD_Female, C_Male |
| II | AUD_Male, C_Female |
| III | One sex remains stable |

**Figure 6.**
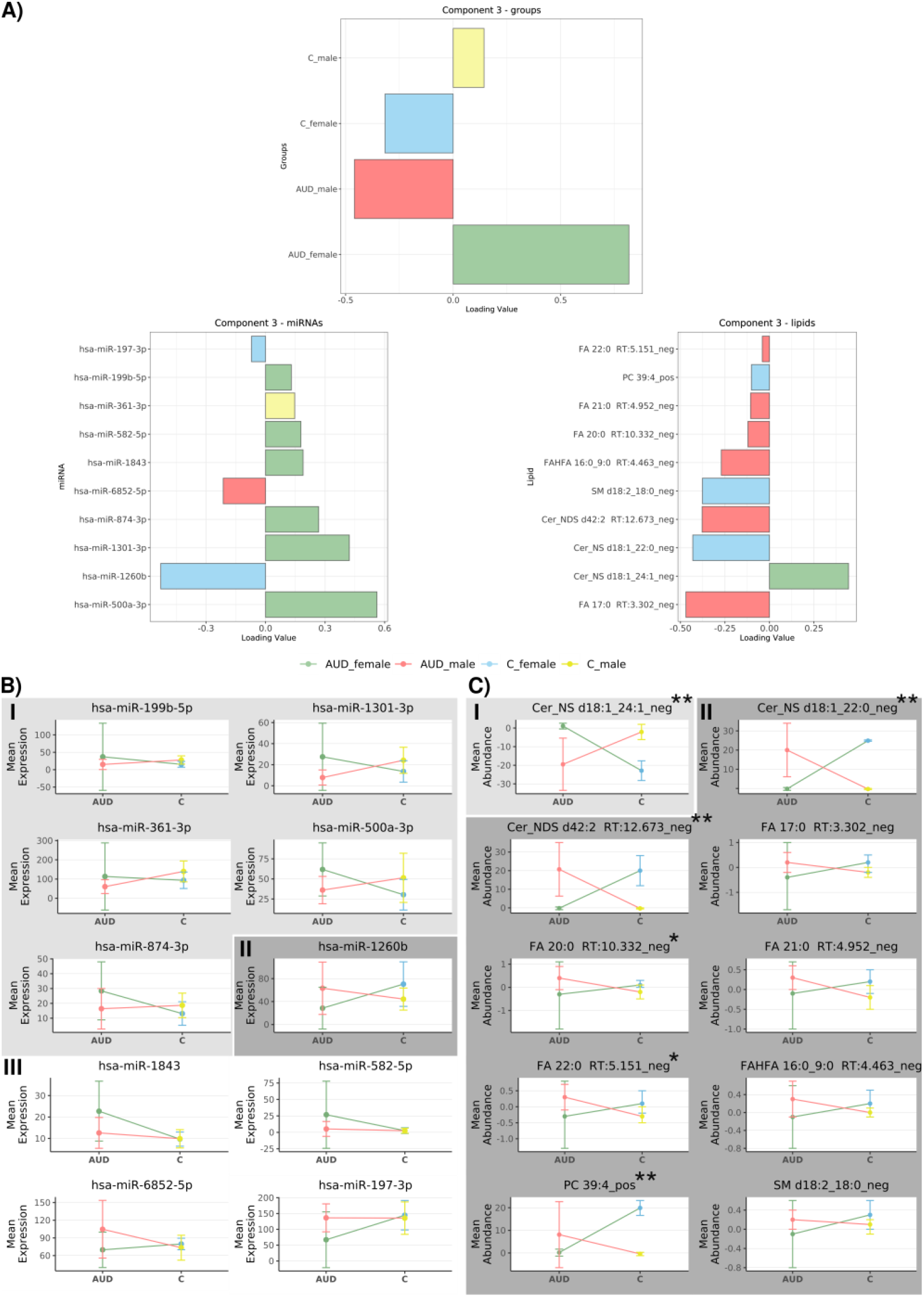
AUD-Sex Signature Reveals Divergent Patterns by Sex and AUD Status. **A)** Loading plot from block.sPLS integration for Component 3. **B)** Mean expression levels of miRNAs and **C)** mean abundance of lipids that explain the variance in Component 3 across experimental groups. *denotes a trend (0.05 < adjusted p-value < 0.055) and **denotes statistical significance (adjusted p-value < 0.01) in the regression analysis (IS comparison of differential expression/abundance analysis). Background colors (light grey, dark grey, and white) indicate the patterns I, II, and III.

This panel includes 40 features (lipids and miRNAs) identified through block.sPLS. Positive (+) or negative (-) symbols indicate the correlation of the feature with the AUD condition. DEA/DAA column indicates the sex where the feature presented a statistical significance in the individual regression analyses. AUD: Alcohol use disorder, DEA: Differential Expression Analysis of miRNome, DAA: Differential Abundance Analysis of lipidome / M: male, F: female.

This panel includes 20 features (lipids and miRNAs) identified through block.sPLS. The pattern indicates the group where the feature is higher in each sex (AUD or Control group). X in the DEA and DAA columns indicates statistical significance (adjusted p-value < 0.01) in the IS comparison in the individual regression analyses (DEA/DAA); # denotes a trend towards significance (0.05 < adjusted p-value < 0.055). *AUD: Alcohol Use Disorder, DEA: Differential Expression Analysis of miRNome, DAA: Differential Abundance Analysis of lipidome, F: female, M: male, IS: Impact of Sex Comparison*.

## Discussion

While lipidomic or miRNomic analyses of EVs have been previously applied to the AUD research [6,35], this study, to the best of our knowledge, represents the first to implement a multi-omics integration of both omic layers, providing a more comprehensive molecular perspective of AUD. The analysis of EVs identified distinct molecular signatures composed of specific combinations of highly correlated miRNAs and lipids. Our multi-omic approach yielded two miRNA-lipid signatures: 1) the AUD signature, comprising a set of 40 biomarkers (20 miRNAs and 20 lipids) that differentiate individuals with AUD from controls; and 2) the AUD-sex signature, consisting of 20 biomarkers (10 miRNAs and 10 lipids) that highlight sex-based differences in the impact of AUD. Our data implicate the integrated features (Tables 1 and 2) in alcohol-related pathophysiological processes and effectively discriminate individuals with AUD from healthy individuals, including sex-specific response to AUD. Importantly, we chose these variables not for their predictive strength, but for their collective contribution to maximizing group separation in the multivariate space when applying the block.sPLS model.

Prior to integration, we performed single-omic univariate differential analyses of lipidomic [6] and miRNomic datasets to identify individual features associated with AUD and sex. These analyses allowed us to explore potential biomarkers with strong individual effects and characterize group- and sexspecific expression patterns in each distinct molecular layer. In the context of miRNomics, we identified 7 (females) and 16 (males) significantly altered miRNAs, with only one miRNA in common. These overlapping features offer additional robustness, suggesting sex-specific biomarker candidates with potentially greater individual effect sizes (e.g., hsa-miR-99b-3p). The RT-qPCR validation confirmed the dysregulation of three key miRNAs in AUD, revealing sexspecific patterns and pathological implications. hsalet-7b-5p was downregulated in AUD males, consistent with prior findings linking it to hippocampal neurodegeneration and addiction [35,36] while ethanol-induced neuroimmune responses involve let-7b/HMGB1 release from microglia [37]. Hsa-miR-194-5p showed upregulation in AUD females, aligning with studies on neuron-derived EVs in heavy drinkers [30] and alcoholic brain tissue [31], and its association with NF-κB signaling suggests roles in inflammation and hepato-carcinoma [38]. Hsa-miR-21-5p was upregulated in both sexes, tied to chronic AUD via IL6R/STAT3-mediated neuroinflammation [39] and liver disease [40], contrasting with acute alcohol exposure studies [32]. These findings highlight the potential of these miRNAs as sex-specific biomarkers and underscore their roles in AUD-related neurodegeneration, immune activation, and organ damage, emphasizing the need to distinguish acute versus chronic alcohol effects in biomarker research.

AUD exerts complex biological effects, increasing the risk of cardiovascular, neurological, metabolic, hepatic, and oncological diseases [41,42]. Although we possess data regarding sex-based differences in AUD, particularly in inflammatory responses [43,44], they are frequently overlooked in research. The historic underrepresentation of females in AUD studies has resulted in critical gaps in understanding, diagnosis, and treatment, which may contribute to poorer clinical outcomes and underscore the urgent need for more inclusive and sex-informed research and therapeutic strategies [45]. In our study, we identified sex-specific miRNomic and lipidomic signatures associated with alcohol-induced molecular alterations, including divergent functional responses to ethanol exposure. Several affected pathways related to inflammation, autophagy, and cellular stress, representing processes commonly implicated in alcohol-related pathologies and cancer. Among the seven cancer-related KEGG pathways identified, we observed the recurrent involvement of crucial genes (e.g., MAP2K1, BRAF, KRAS, MAPK3, ARAF, RAF1, and MAPK1) frequently mutated or dysregulated in cancer that play central roles in tumor progression by promoting proliferation, survival, invasion, and metastasis through activation of the MAPK/ERK signaling cascade [46]. Interestingly, differentially expressed miRNAs predicted to target these oncogenes exhibited marked sex specificity, with only one miRNA shared between males and females. This sex-dependent regulatory pattern may reflect distinct molecular adaptations to chronic alcohol exposure and could contribute to differential cancer susceptibility and progression. Notably, several features from the AUD–sex signature have previously been associated with cancer in other contexts; for example, hsa-miR-1260b has been implicated in pancreatic, lung, ovarian, breast, and myxofibrosarcoma cancers [47–51].

Our data also revealed pronounced sex-specific alterations in autophagy-related regulatory pathways, including those mediated by p53, ubiquitination, Transforming Growth Factor beta (TGF-β), TORC1, the MAPK cascade, mTOR, ErbB signaling, and histone H3 methylation. Prior studies underscored the importance of autophagy in responses to substance-induced stress, particularly alcohol exposure [52–54] and alcohol-associated pathologies such as hepatotoxicity [55]. For instance, alcohol triggers p53-mediated apoptosis in neural crest cells by suppressing TORC1 activity and ribosomal biogenesis, illustrating the complex interplay of these pathways in alcohol-induced neurotoxicity [56]. Chronic alcohol intake also inhibits autophagy and enhances apoptosis in hepatic tissue [57]; moreover, several reports documented sex-based differences in autophagy regulation [58,59], which may help to explain the divergent responses observed in our data in AUD. These differences may contribute to varying susceptibilities to alcohol-induced organ damage and have important implications for the development of sex-informed therapeutic strategies targeting autophagy. These sex-dependent molecular patterns extended to oxidative stress responses as well. Although evidence suggests that males are generally more susceptible to oxidative damage and chronic inflammation, whereas females—particularly before menopause—tend to exhibit stronger inflammatory responses and enhanced antioxidant capacity [60], women are generally more susceptible to developing liver disease or cardiomyopathy after consuming alcohol than men, as well as having a higher incidence of depression and anxiety [61]. Although direct evidence for sex-specific stress granule responses to chronic alcohol exposure are limited, cytoplasmic stress granule formation has been reported in *Saccharomyces cerevisiae* under ethanol stress [62], and sex differences in stress granule composition have been documented in murine models during cold exposure [63]. Our data revealed a divergent expression pattern consistent with sex-specific regulation of stress granules, and one miRNA from our AUD–Sex signature, hsa-miR-197-3p, has been linked to exosomal miRNAs in depression [64] and to circulating miRNAs associated with endothelial dysfunction and cardiometabolic risk in conditions such as Kawasaki disease [65].

Finally, the sphingomyelin biosynthetic process emerged in our sex-differential analysis, reinforcing the notion of sex-specific regulation of sphingolipid metabolism in alcohol-induced brain alterations. This finding agrees with previous reports describing sexual dimorphism in white matter vulnerability and behavioral outcomes in AUD patients [66]. Our previous studies highlighted the role of acid sphingomyelinase in modulating emotional phenotypes and an association with sex-related comorbidities such as depression and anxiety in AUD [6]; moreover, a lipid-driven mechanism involving sphingomyelinases and mitochondria-associated endoplasmic reticulum membranes has been implicated in ethanol-induced EV secretion in glial cells [67]. Our current findings support the hypothesis that sex-specific activity in sphingolipid pathways contributes to divergent neurobiological trajectories in males and females following chronic alcohol exposure. Furthermore, recent evidence has revealed that ceramide production via neutral sphingomyelinase 2 regulates exosomal miRNA secretion and that inhibiting this pathway reduces exosome formation [68,69], a process that also appears to be modulated by sex.

Multi-omic analysis proved particularly valuable for uncovering sex-related differences in disease mechanisms by capturing complex expression patterns that remained undetected in univariate comparisons [66]. While the differential expression analysis of the miRNomic data failed to reveal any significant alterations in the sex-based (IS) comparison at the individual variable level, the integrative multi-omic analysis uncovered a subset of sex-associated miRNAs, underscoring the influence of sex in AUD when analyzing distinct omic layers together. Notably, Component 3 revealed a distinct AUD–sex signature, suggesting that sex-based differences in AUD may not only arise from differences in magnitude but also from qualitatively distinct expression profiles (e.g., upregulation in one sex and stable expression in the other). Additionally, this integrative strategy provided complementary evidence that reinforced findings from the single-omic analysis; specifically, 6 of 10 lipids contributing to Component 3 also displayed significance or consistent trends in the individual lipidomic IS analysis [6]. These lipids belong to classes previously associated with male sex, inflammation, and hepatotoxicity, including ceramides (Cer), sphingomyelins (SM), and fatty acids (FA). Moreover, a substantial proportion of the 40 lipids contributing to AUD- and sex-based separations in Components 1 and 2 overlapped with the most deregulated species identified in the univariate analyses, particularly lysophosphatidylcholines (LPC), phosphatidylcholines (PC), sphingomyelins (SM), and ceramides (Cer).

As emerging disciplines, multi-omics and EV analysis represent the forefront of scientific innovation; however, they still face several limitations. The limited number of plasma samples from AUD females represents a significant limitation of this study; women often do not underreport alcohol-related problems and this issue in women tends to be overlooked in primary care and other settings [70]. Another limitation arises from integrating multi-omics data from the same individuals, which commonly results in the loss of samples due to missing data across omic layers. While newer methodologies can mitigate this problem, the challenge remains a significant limitation in multi-omics-based studies [71]. Although integration enhances class discrimination, this approach does not establish direct links between transcripts and metabolites, which limits the potential for deeper biological interpretation due to the inherent differences between distinct omic layers [72]. Furthermore, the lack of a consensual optimal miRNA normalization strategy highlights the difficulty in comparing studies [73].

In conclusion, this study proposes an AUD signature for predictive modeling, which could contribute to diagnostic applications, and an AUD-sex signature to capture sex-specific molecular biomarkers, enabling a deeper understanding of how biological responses to alcohol differ between females and males. This latter aspect contributes to reducing the persistent gap in female representation within alcohol-related research. Together, our findings support the identification of biologically grounded and sex-informed molecular patterns with potential translational value, which can ultimately be translated into non-invasive biomarker panels to enhance the detection, monitoring, and personalized intervention strategies for AUD.

## Declarations

### Ethics approval and consent to participate

Human plasma samples were used in accordance with the Declaration of Helsinki and were approved by the Ethics Committee of the University Hospital of Salamanca (March 2010). Written informed consent was obtained from each participant.

## Consent for publication

Not applicable

## Availability of data and materials

The normalized and analyzed datasets, along with the programming scripts, are available in the GitHub repository https://github.com/carlapercle/SexEVEthOmics and through the interactive web platform https://carpercle.shinyapps.io/SexEVEthOmics/.

The small RNA-seq data have been deposited in the NCBI Sequence Read Archive (SRA) under accession number PRJNA1291119.

## Authors’ contributions

CPC analyzed the data; MP and FGG designed and supervised the bioinformatics analysis; SK and KL guided in the multi-omics approach; MFR and MM obtained human plasma samples; SM isolated EVs from human plasma; CPC and BMU designed and implemented the web tool; CPC, MP, FGG and CGR wrote the manuscript; SM and CPC designed the graphical abstract; CPC, MP, FGG and CGR helped in the interpretation of the results; CPC, CGR, MM, MP and FGG writing-review and editing; MP and FGG conceived the work. All authors read and approved the final manuscript.

## Acknowledgments

The authors thank the Príncipe Felipe Research Center (CIPF) for providing access to the cluster, which is co-funded by the European Regional Development Fund (FEDER) in the Valencian Community (2014-2020). The authors also thank the Genomics and Genetics Service at the Príncipe Felipe Research Centre and Stuart P. Atkinson for reviewing the manuscript.

