## Supporting information for "Multi-omics integration reveals sex-based differences in the circulating extracellular vesicle lipidome and miRNome of alcohol use disorder patients"

### **Table S1. Cross-omic sample correspondence.** Samples in the same row represent the same individual across different omics. Empty cells indicate no available data for that omic.

| **Lipid Sample** | **miRNA Sample** | **Integration Sample** | **Age** |
| --- | --- | --- | --- |
| AUD_F1 | AUD_Female2 | AUD_female1 | 31 |
| AUD_F2 | AUD_Female3* | - | 41 |
| AUD_F3 | AUD_Female4 | AUD_female2 | 45 |
| AUD_F4 | AUD_Female5 | AUD_female3 | 58 |
| AUD_F5 | - | - | 25 |
| AUD_M1 | - | - | 57 |
| - | AUD_Male2 | - | 41 |
| AUD_M2 | AUD_Male3 | AUD_male1 | 55 |
| AUD_M3 | AUD_Male4 | AUD_male2 | 33 |
| AUD_M4 | AUD_Male5 | AUD_male3 | 51 |
| AUD_M5 | AUD_Male6 | AUD_male4 | 38 |
| AUD_M6 | AUD_Male7 | AUD_male5 | 52 |
| C_F1 | C_Female1 | C_female1 | 42 |
| C_F2 | C_Female2 | C_female2 | 28 |
| C_F3 | C_Female3 | C_female3 | 49 |
| C_F4 | C_Female6 | C_female4 | 30 |
| C_F5 | C_Female4 | C_female5 | 51 |
| - | C_Female5 | **-** | 31 |
| C_F6 | **-** | **-** | 27 |
| C_M1 | C_Male1 | C_male1 | 45 |
| C_M2 | C_Male2 | C_male2 | 43 |
| C_M3 | C_Male3 | C_male3 | 44 |
| C_M4 | C_Male4 | C_male4 | 38 |
| C_M5 | C_Male5 | C_male5 | 52 |
| C_M6 | **-** | **-** | 51 |

*Sample removed during quality control filtering.

### Table S2. Nucleotide sequences of the primers used for qPCR of micro-RNAs.

| **MicroRNA** | **Chromosome**  **location** | **Accession**  **Number (#)** | **Mature primer sequences (5’ to 3’)** |
| --- | --- | --- | --- |
| **hsa-miR-370-3p** | Chr.14: 100911139 - 100911213 [+] on Build GRCh38 | MIMAT0000722 | GCCUGCUGGGGUGGAACCUGGU |
| **hsa-let-7b-5p** | Chr.22: 46113686 - 46113768 [+] on Build GRCh38 | MIMAT0000063 | UGAGGUAGUAGGUUGUGUGGUU |
| **hsa-miR-194-5p** | Chr.1: 220118157 - 220118241 [-] on Build GRCh38 | MIMAT0000460 | UGUAACAGCAACUCCAUGUGGA |

### Table S3. Summary of significant results for Differential Expression Analysis.

|  | **UP** | **DOWN** | **TOTAL** |
| --- | --- | --- | --- |
| **IF** | 4 | 3 | 7 |
| **IM** | 12 | 4 | 16 |
| **IS** | 0 | 0 | 0 |

**IF (AUD Impact in Females):** AUD females vs. control females, **IM (AUD Impact in Males):** AUD males vs. control males, **IS (Impact of Sex in AUD):** Sex-dependent differences in AUD (IF − IM). Significance BH p-value ≤ 0.05.

### Table S4. Significant GSA results for the three comparisons (IF, IM, IS) and their intersections.

|  | **GO** | | | **KEGG** |
| --- | --- | --- | --- | --- |
|  | **BP** | **CC** | **MF** |  |
| **IF** | 127 | 63 | 51 | 35 |
| **IM** | 636 | 223 | 273 | 93 |
| **IS** | 650 | 208 | 232 | 96 |
| **IF & IM & IS**  **LOR < 0 IF**  **LOR > 0 IM** | 66 | 43 | 40 | 20 |
| **IF & IM & IS**  **LOR > 0 IF**  **LOR < 0 IM** | 1 | 1 | 2 | 1 |

The table reports the number of significantly enriched functional terms (Gene Ontology: BP, CC, MF; KEGG pathways) identified in:

- **Individual comparisons:**
  - **IF (AUD Impact in Females):** AUD females vs. control females.
  - **IM (AUD Impact in Males):** AUD males vs. control males.
  - **IS (Impact of Sex in AUD):** Sex-dependent differences in AUD (IF − IM).
- **Intersections of all three comparisons (IF & IM & IS).** These represent terms that are:
  - Simultaneously significant in IF, IM, and IS
  - Further filtered by expression direction:
  - **LOR < 0 in IF & LOR > 0 in IM**: Terms where miRNAs are down-regulated in AUD females but up-regulated in AUD males.
  - **LOR > 0 in IF & LOR < 0 in IM**: Terms where miRNAs are up-regulated in AUD females but down-regulated in AUD males.

*Key observations:*

- The intersection identifies pathways robustly associated with AUD that show sex-opposed expression patterns.
- The dramatic difference in term counts (66 vs 1) reveals most shared pathways show female down-regulation/male up-regulation in AUD.
- Few pathways exhibit the opposite pattern (female up-regulation/male dow-nregulation).

GO (Gene Ontology), BP (Biological Process), CC (Cellular Component), MF (Molecular Function), KEGG pathways (Kyoto Encyclopedia of Genes and Genomes pathways), LOR (log odd ratio; positive = higher in AUD group or female-biased, negative = higher in controls or male-biased).

*(See Materials and Methods and FigureS1 for detailed comparison definitions and statistical thresholds.)*

### **Table S5.** Abbreviation of the different lipid subclasses.

| CAR | Acylcarnitine |
| --- | --- |
| Cer_ADS | Ceramide α-hydroxy fatty acid-dihydrosphingosine |
| Cer_AP | Ceramide α-hydroxy fatty acid-phytosphingosine |
| Cer_AS | Ceramide α-hydroxy fatty acid-sphingosine |
| Cer_NDS | Ceramide non-hydroxy fatty acid-dihydrosphingosine |
| Cer_NP | Ceramide non-hydroxy fatty acid-phytosphingosine |
| Cer_NS | Ceramide non-hydroxy fatty acid-sphingosine |
| Chol. esters | Cholesterol esters |
| DAG | Diacylglycerol |
| FA | Fatty acid |
| FAHFA | Fatty acid ester of hydroxyl fatty acid |
| HexCer_NDS | GlucosylCeramide/HexosylCeramidesnon-hydroxyfatty acid-dihydrosphingosine |
| HexCer_NS | GlucosylCeramide/HexosylCeramidesnon-hydroxyfatty acid-sphingosine |
| LPC | Lyso-phosphatidylcholine |
| LPE | Lyso-phosphatidylethanolamine |
| MGDG | Monogalactosyldiacylglycerol |
| OxPC | Oxidized phosphatidylcholine (OxPC) |
| OxPC-O | Oxidized etherphosphatidylcholine (OxEtherPC) |
| PA | Phosphatidic acid |
| PC | Phosphatidylcholine |
| PC-O | Etherphosphatidylcholine (EtherPC) |
| PE | Phosphatidylethanolamine |
| PE-O | Etherphosphatidylethanolamine (EtherPE) |
| PI | Phosphatidylinositol |
| SHexCer | SulfoglucosylCeramide/SulfohexosylCeramide |
| SM | Sphingomyelin |
| TAG | Triacylglycerol (TG) |

### Table S6. Compilation of disease-associated microRNAs and lipids identified in this study and supported by previous literature.

| **Category** | **microRNA/lipid** | **Condition** | **Sex-differences** | **References** |
| --- | --- | --- | --- | --- |
| Alcohol / TLR4 | hsa-miR-194-5p | AUD |  | [Ref](https://www.nature.com/articles/s41398-024-02874-3#Sec20) |
|  | hsa-miR-148a-3p | AUD |  | [Ref1](https://www.sciencedirect.com/science/article/pii/S000629522100054X), [Ref2](https://pubmed.ncbi.nlm.nih.gov/33895159/), [Ref3](https://pmc.ncbi.nlm.nih.gov/articles/PMC7387083/) |
|  | hsa-miR-21-5p | AUD |  | [Ref1](https://doi.org/10.1186/s13293-017-0158-2), [Ref2](https://doi.org/10.3390/ijms21186730) |
|  | hsa-miR-148a-5p | AUD |  | [Ref](http://ref) |
|  | hsa-miR-486-5p | AUD |  | [Ref](https://doi.org/10.1186/s13293-017-0158-2) |
|  | hsa-miR-146a-5p | Male |  | [Ref](https://doi.org/10.1186/s13293-017-0158-2) |
|  | hsa-miR-16-5p | Male |  | [Ref](https://doi.org/10.1186/s13293-017-0158-2) |
| Cancer | hsa-miR-363-3p | AUD |  | [Ref1](https://www.aging-us.com/article/204398/text), [Ref2](https://pmc.ncbi.nlm.nih.gov/articles/PMC9405730/), [Ref3](https://www.sciencedirect.com/science/article/pii/S0022202X22026409), [Ref4](https://pmc.ncbi.nlm.nih.gov/articles/PMC8798052/) |
|  | hsa-miR-589-5p | AUD |  | [Ref1](https://www.mdpi.com/1422-0067/23/20/12625), [Ref2](https://jeccr.biomedcentral.com/articles/10.1186/s13046-021-01861-6) |
|  | hsa-miR-7706 | AUD |  | [Ref](https://pmc.ncbi.nlm.nih.gov/articles/PMC8798052/) |
|  | hsa-miR-194-5p | AUD |  | [Ref1](https://doi.org/10.1038/s41556-021-00805-8), [Ref2](https://cancerci.biomedcentral.com/articles/10.1186/s12935-022-02835-0), [Ref3](https://pmc.ncbi.nlm.nih.gov/articles/PMC7104536/#Sec2) |
|  | hsa-miR-148a-3p | AUD |  | [Ref1](https://pmc.ncbi.nlm.nih.gov/articles/PMC8990464/), [Ref2](https://europepmc.org/article/pmc/7817948#s4) |
|  | hsa-miR-21-5p | AUD |  | [Ref1](https://www.mdpi.com/1422-0067/23/20/12625), [Ref2](https://www.sciencedirect.com/science/article/pii/S0022202X22026409), [Ref3](https://pmc.ncbi.nlm.nih.gov/articles/PMC8798052/) |
|  | hsa-miR-148a-5p | AUD |  | [Ref](https://pmc.ncbi.nlm.nih.gov/articles/PMC8798052/) |
|  | hsa-miR-130b-3p | AUD |  | [Ref1](https://www.sciencedirect.com/science/article/pii/S0022202X22026409), [Ref2](https://doi.org/10.1038/s41556-021-00805-8) |
|  | hsa-miR-223-5p | AUD |  | [Ref](https://pmc.ncbi.nlm.nih.gov/articles/PMC8798052/) |
|  | hsa-miR-99b-3p | AUD |  | [Ref](https://pmc.ncbi.nlm.nih.gov/articles/PMC8798052/) |
|  | hsa-miR-192-5p | AUD |  | [Ref](https://www.frontiersin.org/journals/pharmacology/articles/10.3389/fphar.2021.614068/full) |
|  | hsa-miR-486-5p | AUD |  | [Ref](https://pmc.ncbi.nlm.nih.gov/articles/PMC8798052/) |
|  | hsa-miR-197-3p | Interaction | ♀ < ♂ | [Ref1](https://pmc.ncbi.nlm.nih.gov/articles/PMC6341871/), [Ref2](https://pmc.ncbi.nlm.nih.gov/articles/PMC9917813/) |
|  | hsa-miR-1260b | Interaction | ♀ < ♂ | [Ref1](https://pmc.ncbi.nlm.nih.gov/articles/PMC9207358/), [Ref2](https://ovarianresearch.biomedcentral.com/articles/10.1186/s13048-021-00878-x), [Ref3](https://www.nature.com/articles/s41419-021-04024-9#Sec23) |
|  | hsa-miR-199b-5p | Interaction | ♀ > ♂ | [Ref](https://pmc.ncbi.nlm.nih.gov/articles/PMC6144876/) |
|  | hsa-miR-582-5p | Interaction | ♀ > ♂ | [Ref1](https://www.turkishneurosurgery.org.tr/pdf/pdf_JTN_2691.pdf), [Ref2](https://pmc.ncbi.nlm.nih.gov/articles/PMC8798052/) |
|  | hsa-miR-361-3p | Interaction | ♀ > ♂ | [Ref](https://pmc.ncbi.nlm.nih.gov/articles/PMC8798052/) |
|  | hsa-miR-146a-5p | Male |  | [Ref1](https://www.mdpi.com/1422-0067/23/20/12625), [Ref2](https://www.sciencedirect.com/science/article/pii/S0022202X22026409) |
|  | hsa-miR-24-3p | Male |  | [Ref](https://pmc.ncbi.nlm.nih.gov/articles/PMC8798052/) |
|  | hsa-miR-424-3p | Male |  | [Ref](https://pmc.ncbi.nlm.nih.gov/articles/PMC8798052/) |
|  | hsa-miR-1294 | Male |  | [Ref](https://pmc.ncbi.nlm.nih.gov/articles/PMC8798052/) |
|  | hsa-miR-152-3p | Male |  | [Ref](https://pmc.ncbi.nlm.nih.gov/articles/PMC8798052/) |
|  | hsa-miR-145-3p | Male |  | [Ref1](https://www.mdpi.com/2072-6694/13/13/3287), [Ref2](https://pmc.ncbi.nlm.nih.gov/articles/PMC8798052/) |
|  | hsa-miR-1271-5p | Male |  | [Ref](https://pmc.ncbi.nlm.nih.gov/articles/PMC8798052/) |
|  | hsa-miR-425-3p | Male |  | [Ref](https://pmc.ncbi.nlm.nih.gov/articles/PMC8798052/) |
|  | hsa-miR-454-3p | C |  | [Ref1](https://doi.org/10.1158/1535-7163.mct-18-0725), [Ref2](https://pmc.ncbi.nlm.nih.gov/articles/PMC8020204/), [Ref3](https://ascopubs.org/doi/10.1200/JCO.2019.37.15_suppl.e15108) |
|  | hsa-let-7b-5p | C |  | [Ref](https://pmc.ncbi.nlm.nih.gov/articles/PMC8798052/) |
|  | hsa-miR-148b-3p | C |  | [Ref1](https://pmc.ncbi.nlm.nih.gov/articles/PMC6817568/), [Ref2](https://pmc.ncbi.nlm.nih.gov/articles/PMC10234784/), [Ref3](https://www.oncotarget.com/article/2014/text/) |
|  | hsa-miR-331-3p | C |  | [Ref](https://pubmed.ncbi.nlm.nih.gov/30228315/) |
|  | hsa-miR-181d-5p | C |  | [Ref](https://pmc.ncbi.nlm.nih.gov/articles/PMC8798052/) |
|  | hsa-miR-548e-5p | Female |  | [Ref](https://pmc.ncbi.nlm.nih.gov/articles/PMC6485292/#B14) |
|  | hsa-miR-139-5p | Female |  | [Ref](https://pmc.ncbi.nlm.nih.gov/articles/PMC8798052/) |
|  | hsa-miR-299-3p | Female |  | [Ref](https://www.researchgate.net/publication/325179688_Prognostic_value_of_hsa-mir-299_and_hsa-mir-7706_in_hepatocellular_carcinoma) |
|  | hsa-miR-144-3p | Male |  | [Ref](https://pmc.ncbi.nlm.nih.gov/articles/PMC8798052/) |
|  | hsa-miR-30d-5p | Male |  | [Ref](https://pmc.ncbi.nlm.nih.gov/articles/PMC8829117/table/T1/) |
|  | Cer_NDS d41:2 | AUD |  | [Ref](https://www.metabolomicsworkbench.org/data/metstat_studies.php?refmet_name=Cer+41%3A2%3BO2) |
|  | LPE 18:0 | AUD |  | [Ref](https://www.metabolomicsworkbench.org/data/metstat_studies.php?refmet_name=LPE+18%3A0) |
| Cardiovascular | hsa-miR-589-5p | AUD |  | [Ref1](https://www.nature.com/articles/s41598-018-27078-w#Sec13), [Ref2](https://doi.org/10.1111/cts.13307) |
|  | hsa-miR-192-5p | AUD |  | [Ref](https://www.frontiersin.org/journals/pharmacology/articles/10.3389/fphar.2021.614068/full) |
|  | hsa-miR-197-3p | Interaction | ♀ < ♂ | [Ref](https://doi.org/10.1186/s12864-017-3533-9) |
|  | hsa-miR-16-5p | Male |  | [Ref](https://doi.org/10.1002/iub.2189) |
|  | LPE 18:0 | AUD |  | [Ref](https://www.metabolomicsworkbench.org/data/metstat_studies.php?refmet_name=LPE+18%3A0) |
|  | SM d44:2 | AUD |  | [Ref](https://www.metabolomicsworkbench.org/data/metstat_studies.php?refmet_name=SM+44%3A2%3BO2) |
| Inflammatory / Immune | hsa-miR-589-5p | AUD |  | [Ref](https://pmc.ncbi.nlm.nih.gov/articles/PMC10722155/table/T1/#B20) |
|  | hsa-miR-194-5p | AUD |  | [Ref](https://doi.org/10.1007/s10735-025-10464-w) |
|  | hsa-miR-148a-3p | AUD |  | [Ref1](https://pmc.ncbi.nlm.nih.gov/articles/PMC10380621/), [Ref2](https://www.ncbi.nlm.nih.gov/pmc/articles/PMC3912347/) |
|  | hsa-miR-21-5p | AUD |  | [Ref1](https://pmc.ncbi.nlm.nih.gov/articles/PMC9800721/), [Ref2](https://pmc.ncbi.nlm.nih.gov/articles/PMC3505968/), [Ref3](https://pmc.ncbi.nlm.nih.gov/articles/PMC10722155/table/T1/#B20), [Ref4](https://onlinelibrary.wiley.com/doi/10.1111/liv.15682) |
|  | hsa-miR-148a-5p | AUD |  | [Ref](https://pubmed.ncbi.nlm.nih.gov/24175965/) |
|  | hsa-miR-130b-3p | AUD |  | [Ref](https://pubmed.ncbi.nlm.nih.gov/25653823/) |
|  | hsa-miR-192-5p | AUD |  | [Ref](https://www.frontiersin.org/journals/pharmacology/articles/10.3389/fphar.2021.614068/full) |
|  | hsa-miR-146a-5p | Male |  | [Ref](https://www.frontiersin.org/journals/pharmacology/articles/10.3389/fphar.2021.614068/full) |
|  | hsa-miR-144-3p | Male |  | [Ref](https://www.elsevier.es/en-revista-clinics-22-articulo-characterization-microrna-profile-in-rheumatoid-S1807593224001182) |
|  | hsa-let-7b-5p | C |  | [Ref](https://www.frontiersin.org/journals/pharmacology/articles/10.3389/fphar.2021.614068/full) |
|  | hsa-miR-331-3p | C |  | [Ref](https://doi.org/10.1007/s00280-020-04122-z) |
|  | hsa-miR-548e-5p | Female |  | [Ref](https://pubmed.ncbi.nlm.nih.gov/31588496/) |
|  | hsa-miR-145-3p | Male |  | [Ref](https://doi.org/10.2147/cmar.s482688) |
|  | hsa-miR-199b-5p | Interaction | ♀ > ♂ | [Ref](https://elifesciences.org/reviewed-preprints/92645) |
|  | LPE 18:0 | AUD |  | [Ref](https://www.metabolomicsworkbench.org/data/metstat_studies.php?refmet_name=LPE+18%3A0) |
| Metabolic | hsa-miR-589-5p | AUD |  | [Ref](https://pmc.ncbi.nlm.nih.gov/articles/PMC6586838/) |
|  | hsa-miR-148a-3p | AUD |  | [Ref](https://pmc.ncbi.nlm.nih.gov/articles/PMC6586838/) |
|  | hsa-miR-192-5p | AUD |  | [Ref](https://www.frontiersin.org/journals/pharmacology/articles/10.3389/fphar.2021.614068/full) |
|  | hsa-miR-21-5p | AUD |  | [Ref1](https://cardiab.biomedcentral.com/articles/10.1186/s12933-023-01988-0), [Ref2](https://onlinelibrary.wiley.com/doi/10.1111/liv.15682) |
|  | hsa-miR-664a-3p | Male |  | [Ref](https://journals.plos.org/plosone/article?id=10.1371/journal.pone.0079697) |
| Neurodegenerative / Neurological | hsa-miR-589-5p | AUD |  | [Ref](https://pmc.ncbi.nlm.nih.gov/articles/PMC9405730/) |
|  | hsa-miR-192-5p | AUD |  | [Ref](https://www.frontiersin.org/journals/pharmacology/articles/10.3389/fphar.2021.614068/full) |
|  | hsa-miR-146a-5p | Male |  | [Ref](https://link.springer.com/article/10.1007/s12031-022-02001-1) |
|  | hsa-miR-152-3p | Male |  | [Ref](https://pmc.ncbi.nlm.nih.gov/articles/PMC4177314/#S11) |
|  | Cer_NDS d41:2 | AUD |  | [Ref](https://www.metabolomicsworkbench.org/data/metstat_studies.php?refmet_name=Cer+41%3A2%3BO2) |
|  | LPE 18:0 | AUD |  | [Ref](https://www.metabolomicsworkbench.org/data/metstat_studies.php?refmet_name=LPE+18%3A0) |
| Other | hsa-miR-363-3p | AUD |  | [Ref](https://www.aging-us.com/article/204398/text) |
|  | hsa-miR-194-5p | AUD |  | [Ref](https://www.sciencedirect.com/science/article/pii/S1526590024005571#fig0015) |
|  | hsa-miR-7706 | AUD |  | [Ref1](https://www.sciencedirect.com/science/article/pii/S0888754321001208#f0030), [Ref2](https://www.spandidos-publications.com/10.3892/etm.2021.10149) |
|  | hsa-miR-192-5p | AUD |  | [Ref](https://www.frontiersin.org/journals/pharmacology/articles/10.3389/fphar.2021.614068/full) |
|  | hsa-miR-10399-5p | AUD |  | [Ref](https://www.aging-us.com/article/204398/text) |
|  | hsa-miR-556-5p | C |  | [Ref](https://journals.plos.org/plosone/article?id=10.1371/journal.pone.0216400) |
| Psychiatric disorders | hsa-miR-589-5p | AUD |  | [Ref](https://www.elsevier.es/es-revista-european-journal-psychiatry-431-articulo-differential-exosomal-microrna-profile-in-S0213616317300745) |
|  | hsa-miR-21-5p | AUD |  | [Ref](https://www.ncbi.nlm.nih.gov/pmc/articles/PMC10450008/) |
|  | hsa-miR-192-5p | AUD |  | [Ref](https://www.frontiersin.org/journals/pharmacology/articles/10.3389/fphar.2021.614068/full) |
|  | hsa-miR-197-3p | Interaction | ♀ < ♂ | [Ref](https://www.elsevier.es/es-revista-european-journal-psychiatry-431-articulo-differential-exosomal-microrna-profile-in-S0213616317300745) |
|  | hsa-miR-145-3p | Male |  | [Ref](https://www.elsevier.es/es-revista-european-journal-psychiatry-431-articulo-differential-exosomal-microrna-profile-in-S0213616317300745) |
|  | hsa-miR-144-3p | Male |  | [Ref](https://www.elsevier.es/es-revista-european-journal-psychiatry-431-articulo-differential-exosomal-microrna-profile-in-S0213616317300745) |
|  | hsa-miR-139-5p | Female |  | [Ref](https://www.elsevier.es/es-revista-european-journal-psychiatry-431-articulo-differential-exosomal-microrna-profile-in-S0213616317300745) |

*Category*: Disease group or biological context associated with the listed microRNAs or lipids based on previous literature. *Condition*: Experimental group in which the feature was positively correlated (AUD: alcohol use disorder; C: control; Interaction: feature associated with Component 3). *Sex-differences*: For features correlated with Component 3, this column indicates which sex in the AUD group showed higher expression.

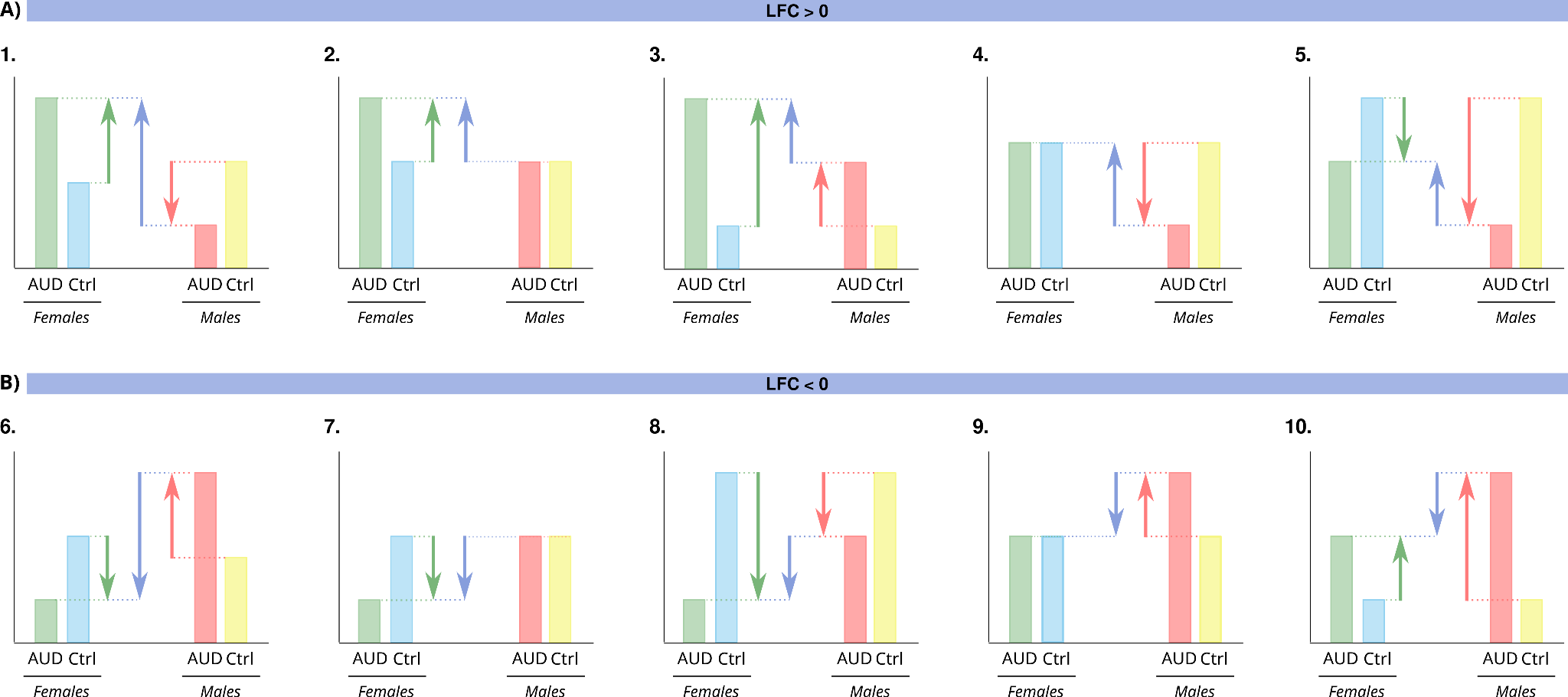

### Figure S1. Bar chart representing the possible causes of a positive (A) and negative (B) LFC/LOR in the IS comparison, depending on the IF or IM comparison.

**Interpretation of functional enrichment results (Figures S2–S5)**

Dot plots display significant functional terms (GO biological processes, cellular components, molecular functions, and KEGG pathways) from gene set analysis (GSA) that are significant in both comparisons IF and IM.

**Comparisons shown:**

1. **Sex-specific AUD effects:**
   - **IF (AUD Impact in Females)**: AUD females vs. control females.
   - **IM (AUD Impact in Males)**: AUD males vs. control males.
2. **Sex-dependent modulation in AUD (IS)**: IF − IM (highlighting terms where AUD effects differ by sex).
3. **Overall AUD impact (C)**: AUD vs. controls.

**Key patterns:**

- **Sex-opposed responses**: Terms significant in both IF and IM with opposite LOR directions (e.g., down-regulated in females but up-regulated in males) are validated in IS. The intersection of IF, IM, and IS identifies robust sex-divergent pathways.
- **Consistent trends**: Most shared terms exhibit female down-regulation / male up-regulation in AUD. Few pathways show the inverse pattern (female up-regulation / male down-regulation).
- **Sex stratification relevance**: Comparison with **C** reveals whether sex-independent analyses mask sexually dimorphic effects.

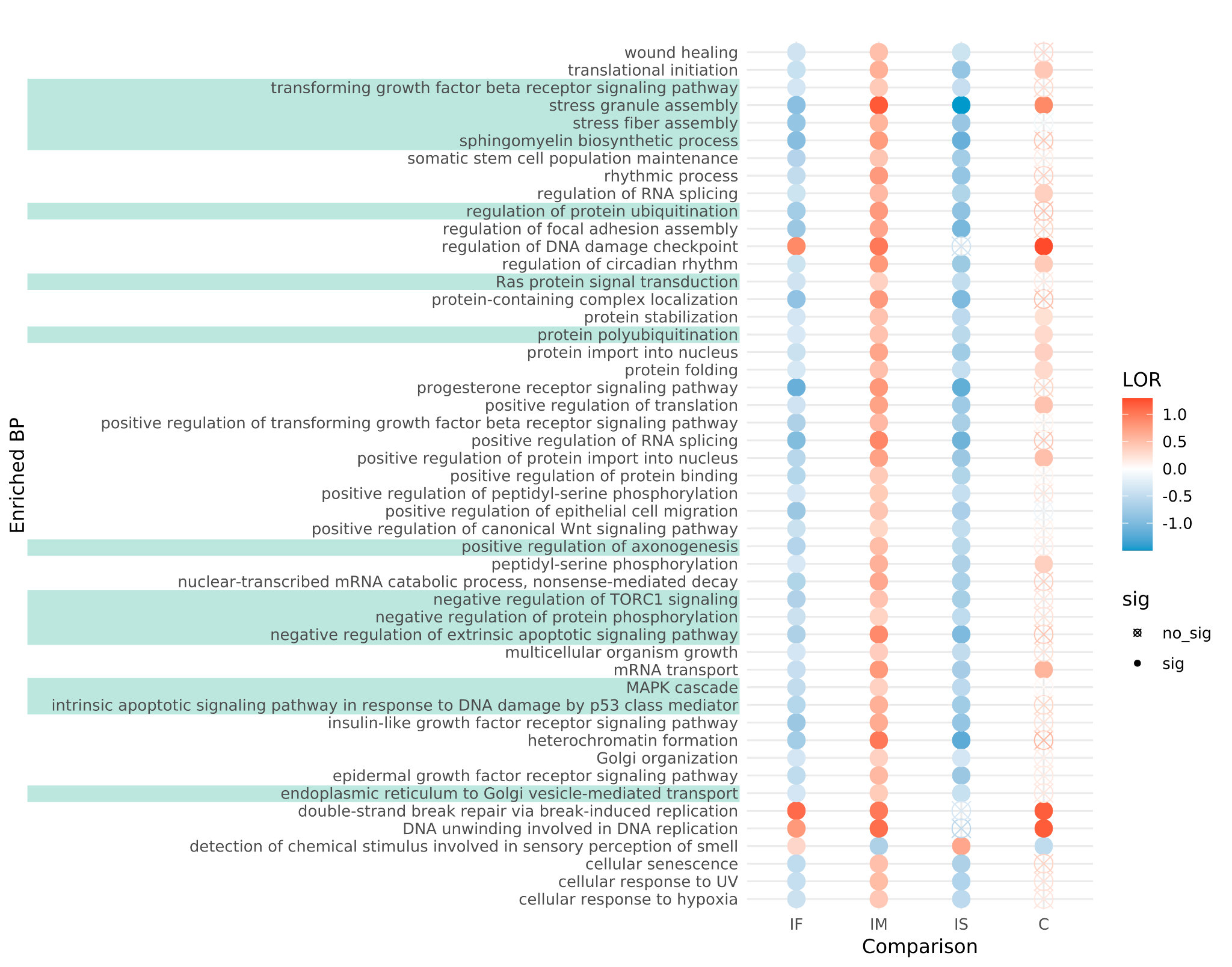

### **Figure S2. Gene Set Analysis (GSA) results showing enriched biological processes (BP) across different comparisons.** Each row represents a significantly enriched BP term, with color intensity indicating the log odds ratio (LOR) of enrichment. Red denotes positive LOR values (overrepresentation in the first group of the comparison), while blue represents negative LOR values (overrepresentation in the second group). Comparisons include AUD Impact in Females (IF), AUD Impact in Males (IM), Impact of Sex in AUD (IS), and the overall comparison (C). Filled circles indicate statistically significant terms, while empty circles denote non-significant results. Highlighted terms (shaded rows) correspond to key BP relevant to AUD.

### **Figure S3. Gene Set Analysis (GSA) results showing enriched cellular components (CC) across different comparisons.** Each row represents a significantly enriched CC term, with color intensity indicating the log odds ratio (LOR) of enrichment. Red denotes positive LOR values (overrepresentation in the first group of the comparison), while blue represents negative LOR values (overrepresentation in the second group). Comparisons include AUD Impact in Females (IF), AUD Impact in Males (IM), Impact of Sex in AUD (IS), and the overall comparison (C). Filled circles indicate statistically significant terms, while empty circles denote non-significant results. Highlighted terms (shaded rows) correspond to key CC relevant to AUD.

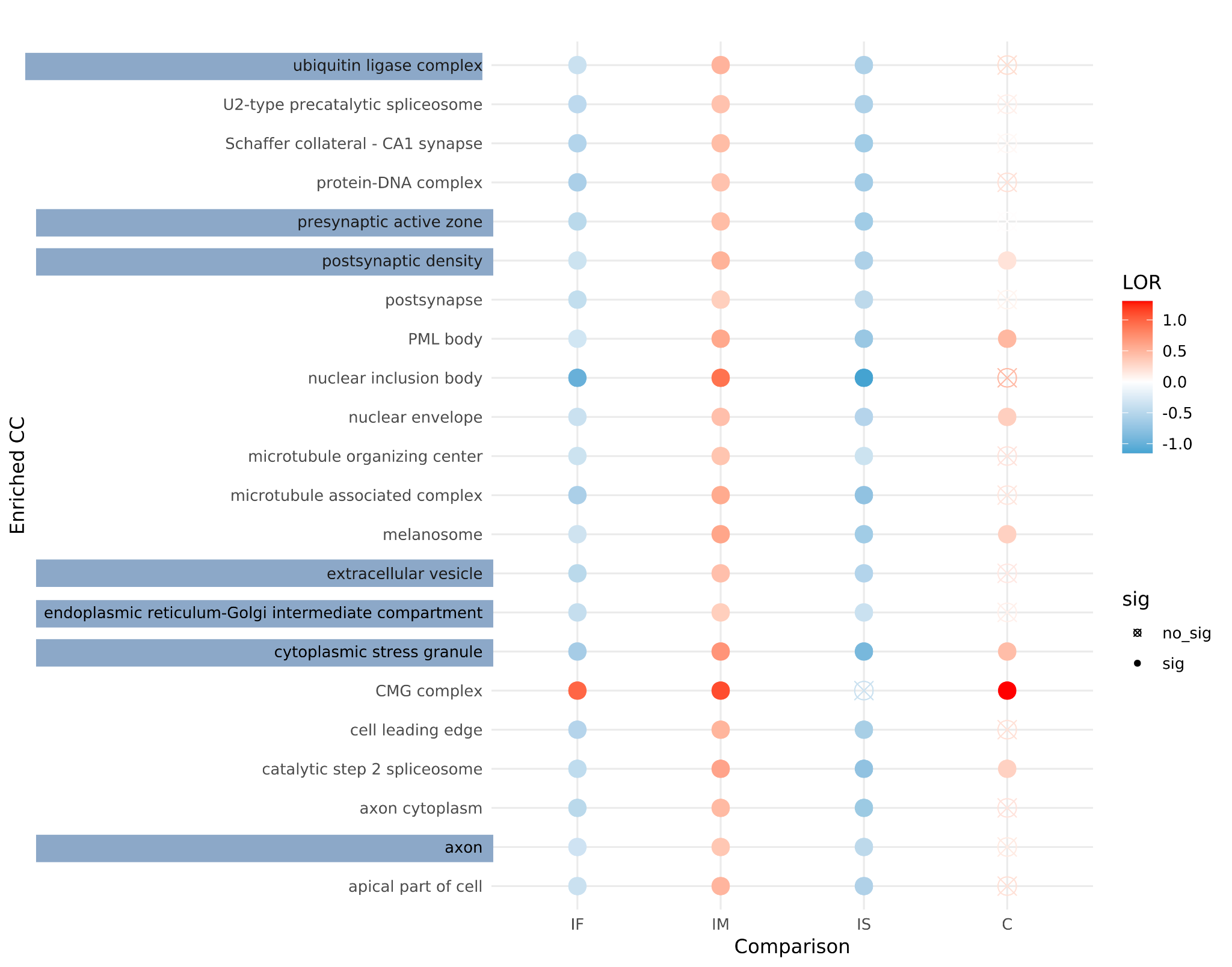

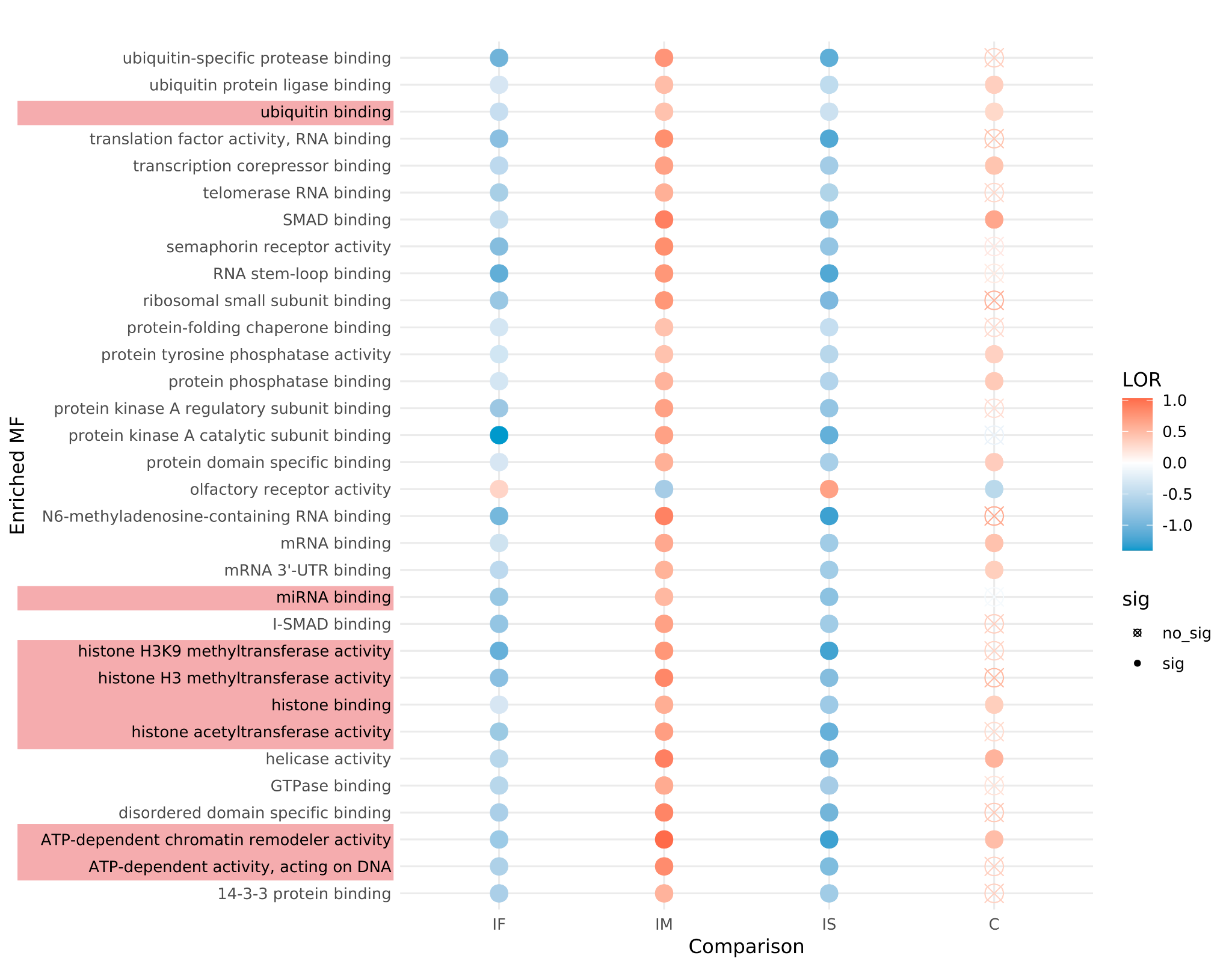

### **Figure S4. Gene Set Analysis (GSA) results showing enriched molecular functions (MF) across different comparisons.** Each row represents a significantly enriched MF term, with color intensity indicating the log odds ratio (LOR) of enrichment. Red denotes positive LOR values (overrepresentation in the first group of the comparison), while blue represents negative LOR values (overrepresentation in the second group). Comparisons include AUD Impact in Females (IF), AUD Impact in Males (IM), Impact of Sex in AUD (IS), and the overall comparison (C). Filled circles indicate statistically significant terms, while empty circles denote non-significant results. Highlighted terms (shaded rows) correspond to key MF relevant to AUD.

### **Figure S5. Gene Set Analysis (GSA) results showing enriched KEGG pathways across different comparisons.** Each row represents a significantly enriched KEGG pathway, with color intensity indicating the log odds ratio (LOR) of enrichment. Red denotes positive LOR values (overrepresentation in the first group of the comparison), while blue represents negative LOR values (overrepresentation in the second group). Comparisons include AUD Impact in Females (IF), AUD Impact in Males (IM), Impact of Sex in AUD (IS), and the overall comparison (C). Filled circles indicate statistically significant terms, while empty circles denote non-significant results.

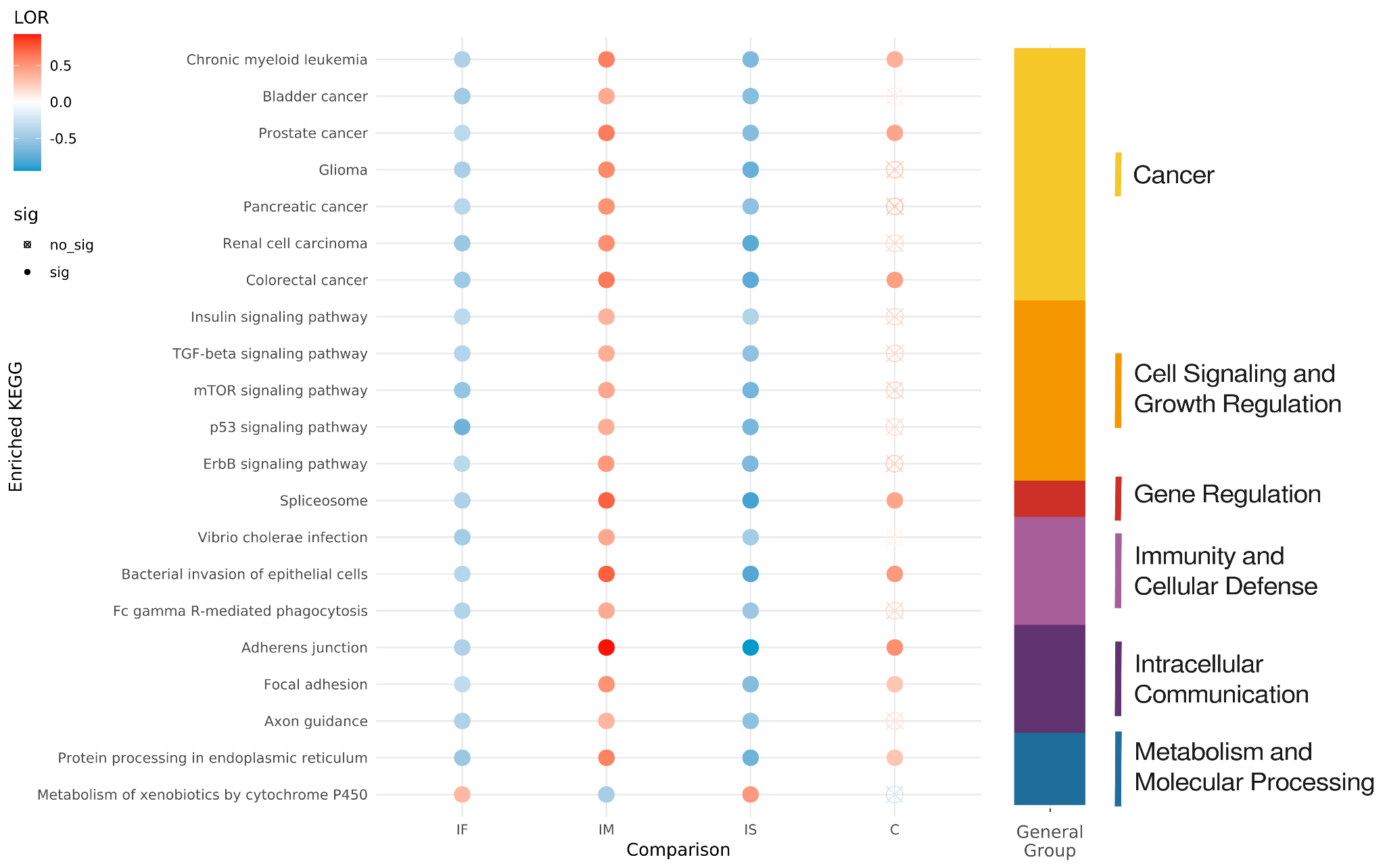

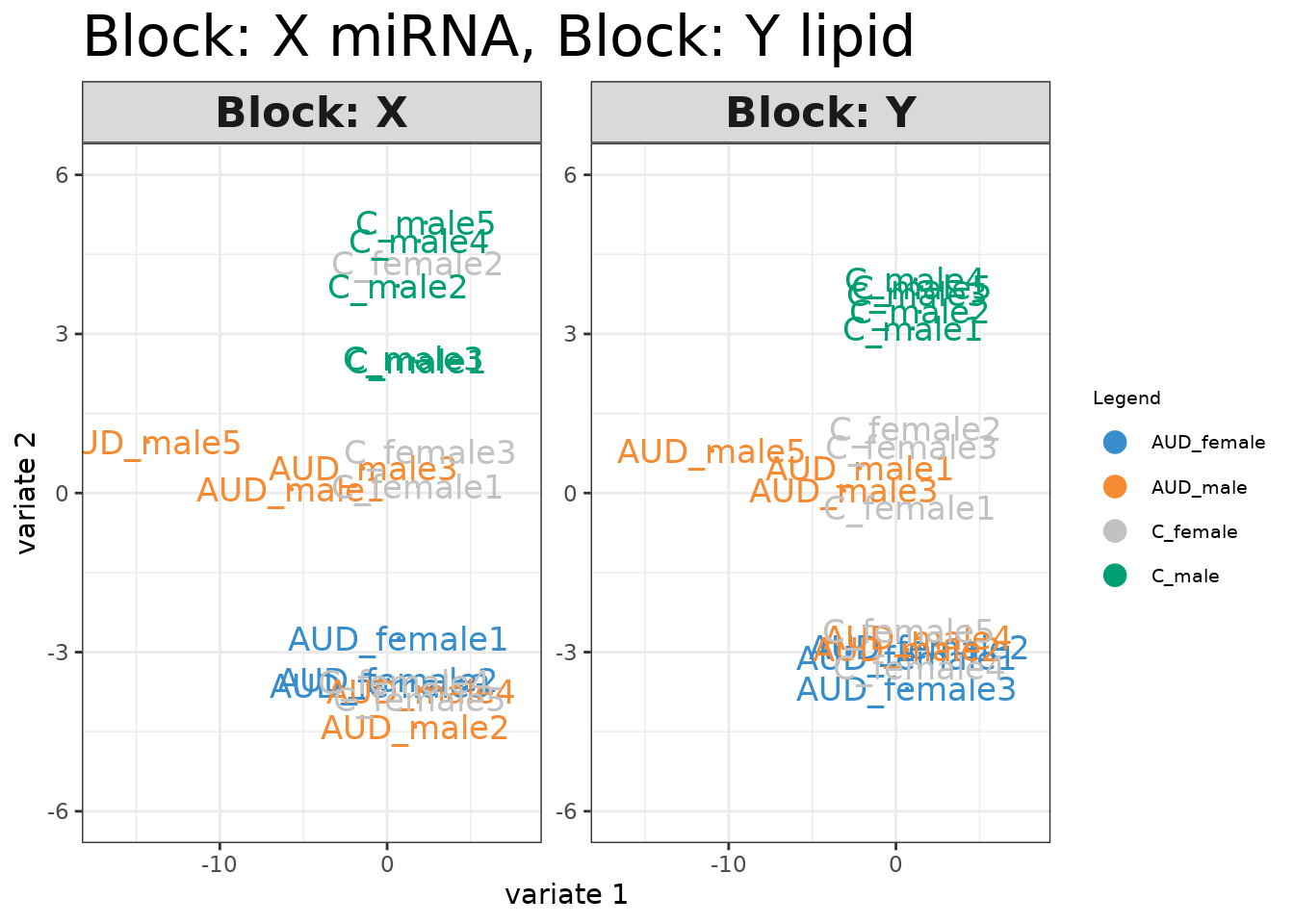

### **Figure S6. Sparse Partial Least Squares (sPLS) analysis integrating miRNA (Block X) and lipid (Block Y) data.** The plot displays sample distributions based on the first two variates, with colors representing the four experimental groups: AUD females (blue), AUD males (orange), control females (gray), and control males (green). The separation of groups reflects the contribution of miRNA and lipid features in distinguishing between conditions.
